# De novo lipid synthesis and polarized prenylation drives cell invasion through basement membrane

**DOI:** 10.1101/2024.02.19.581070

**Authors:** Kieop Park, Aastha Garde, Siddharthan B. Thendral, Adam W. J. Soh, Qiuyi Chi, David R. Sherwood

## Abstract

**Summary:** Invasive cells form large, specialized protrusions to break through basement membrane (BM) matrix barriers. Park et al., reveal a crucial requirement for de novo lipid synthesis and a dynamic polarizing prenylation system to rapidly construct invasive protrusions that breach BMs.

To breach basement membrane (BM), cells in development and cancer use large, transient, specialized lipid-rich membrane protrusions. Using live imaging, endogenous protein tagging, and cell-specific RNAi during *C. elegans* anchor cell (AC) invasion, we demonstrate that the lipogenic SREBP transcription factor SBP-1 drives expression of the fatty acid synthesis enzymes POD-2 and FASN-1 prior to invasion. We show that phospholipid producing LPIN-1 and sphingomyelin synthase SMS-1, which use fatty acids as substrates, produce lysosome stores that build the AC’s invasive protrusion, and that SMS-1 also promotes protrusion localization of the lipid raft partitioning ZMP-1 matrix metalloproteinase. Finally, we discover that the endoplasmic reticulum (ER)-associated HMG-CoA reductase HMGR-1, which generates isoprenoids for prenylation, enriches at the AC invasive front, and that the final ER prenylation enzyme, ICMT-1, localizes to ER exit sites that dynamically polarize to deliver prenylated GTPases for protrusion formation. Together, these results reveal a collaboration between lipogenesis and a polarized lipid prenylation system that drives invasive protrusion formation.

## Introduction

During animal development many cells migrate to form organs and tissues (Aman and Piotrowski, 2010; Scarpa and Mayor, 2016). For example, neural crest undergo an epithelial-to-mesenchymal transition (EMT), detach from the neural tube and travel throughout the embryo to form cartilage, bones, muscles, neurons, and epidermis (Szabo and Mayor, 2018). During their embryonic migrations, cells often traverse basement membrane (BM) —a dense, covalently cross-linked type IV collagen and laminin rich extracellular matrix that enwraps and separates most tissues (Gros and Tabin, 2014; Jayadev and Sherwood, 2017; Kelley et al., 2014; Leonard and Taneyhill, 2020; Moser et al., 2018). Migrating immune cells also transmigrate BM barriers to reach sites of injury and infection in adult organisms (Bahr et al., 2022). Cell invasive behavior is mis-regulated in many diseases, most notably cancer, where metastatic tumor cells hijack normal invasive cellular programs (Paterson and Courtneidge, 2018). To breach BMs, invasive cells use small, matrix metalloproteinase (MMP)-enriched, F-actin-driven plasma membrane protrusions, termed invadosomes (Cambi and Chavrier, 2021). Because of the difficulty of visualizing rare and often stochastic cell invasion events in vivo, invadosomes have been studied most extensively in vitro on 2D surfaces that mimic the planar BM structure (Clarke et al., 2024). The events following BM breaching are less clear due to the challenge of recapitulating invasion in 3D settings. However, ex vivo invasion assays, 3D spheroid cell cultures, and tumor sections have revealed that invadosomes transition into a large, transient protrusion that clears a path through the BM to allow BM transmigration (Hotary et al., 2006; Leong et al., 2014; Nazari et al., 2023; Schoumacher et al., 2010). Although crucial for invasion, the mechanisms regulating this invasive structure remain poorly understood.

Invasive membrane protrusions are large and have the specialized function of BM removal and it is unknown if lipid production or specialized lipid modification systems are required for their formation. Lipid metabolism is complex and involves lipid biosynthesis, external lipid import, lipid storage, and lipid catabolism for energy production (Snaebjornsson et al., 2020). Lipid metabolism can be regulated both at the transcriptional level, often by the sterol regulatory element-binding protein (SREBP) family of transcription factors or by external cues (Röhrig and Schulze, 2016). Exacerbated lipid synthesis is strongly associated with metastatic cancers (Bian et al., 2021; Chen et al., 2018; Lu et al., 2022; Martin-Perez et al., 2022; Vasseur and Guillaumond, 2022). For example, SREBP transcription factors and their key lipogenic enzyme targets are overexpressed in many metastatic cancers and promote invasion in vitro through solubilized non-crosslinked BM extracts. These SREBP targets include acetyl-CoA carboxylase (ACC), which catalyzes the rate limiting step in fatty acid synthesis, fatty acid synthase (FASN), which directs palmitate synthesis, and HMG-CoA reductase (HMGCR), which is the rate limiting enzyme of the mevalonate pathway and necessary for the catalysis of isoprenoids for cholesterol and protein prenylation (Bao et al., 2016; Bian et al., 2021; Gao et al., 2019; Li et al., 2014; Lu et al., 2022; Wang et al., 2022; Xu et al., 2020). It is unclear, however, if these lipogenesis enzymes are required for cells to breach the dense and highly cross-linked BM found in vivo and unknown how lipogenesis might function to promote invasion. Understanding the role of lipogenesis in cell invasion is important, as many lipid synthesis enzyme inhibitors exist that could be strategically used to target invasive behavior (Bian et al., 2021; Broadfield et al., 2021; Vasseur and Guillaumond, 2022).

The anchor cell (AC) is a specialized uterine cell that invades through the BM separating the uterine and vulval tissue in *C. elegans* to initiate uterine-vulval connection (Sherwood and Sternberg, 2003). AC invasion is highly stereotyped, accessible to live imaging, and allows targeted gene knockdown (Kenny-Ganzert and Sherwood, 2023). Like cancer cells, the AC uses dynamic invadosomes to breach the BM (Hagedorn et al., 2013). Following BM breaching, the netrin receptor UNC-40 (vertebrate DCC) traffics to the small hole in the BM and directs lysosome exocytosis to form a large protrusion that clears a path through the BM (Hagedorn et al., 2013; Naegeli et al., 2017). The invasive protrusion is enriched with the glycosylphosphatidylinositol (GPI)-anchored matrix metalloproteinase (MMP) ZMP-1, which helps degrade the BM (Kelley et al., 2019). The UNC-40 receptor and GPI-anchored proteins like ZMP-1 are partitioned to sphingolipid enriched lipid rafts (Herincs et al., 2005; Sangiorgio et al., 2004). In addition, *C. elegans* Rac and Ras-like GTPases that promote F-actin formation for invasive protrusion outgrowth are anchored and enriched in the invasive protrusion through C-terminal lipid prenylation (Costa et al., 2023; Hagedorn et al., 2013; Lohmer et al., 2016; Wang et al., 2014b). Previous work examining energy sources that fuel invasion revealed that the AC does not contain lipid stores nor require lipid import transporters (Garde et al., 2022; Zechner et al., 2017). Whether AC invasion depends on de novo lipid synthesis and elevated GTPase lipid anchoring to form the large specialized invasive protrusion, however, is unknown.

Using live cell imaging, endogenous fluorescently tagged proteins, and AC-specific RNAi, we show that the lipogenic *C. elegans* SREBP transcription factor SBP-1 is required for invasion and invasive protrusion formation. SBP-1 traffics to the AC nucleus prior to invasion and drives elevated expression of genes encoding the fatty acid synthesis enzymes POD-2 (vertebrate acetyl-CoA carboxylase, ACC) and FASN-1 (vertebrate FASN). We show that phospholipid producing phosphatidic acid phosphatase gene *lpin-1* and the sphingomyelin synthase *sms-1* whose protein products use fatty acids made by POD-1 and FASN-1, are upregulated in the AC and required for protrusion formation—LPIN-1 enhances lysosome production required for protrusion formation, and SMS-1 also increases lysosome stores as well as promotes UNC-40 (DCC receptor) and ZMP-1 (MMP) localization within the protrusion.

Finally, we discover that the endoplasmic reticulum (ER)-localized HMGR-1 (vertebrate HMGCR, mevalonate pathway), is enriched in ER at the AC invasive front. HMGR-1 generates isoprenoids for prenylation, and we find that the ER-localized isoprenylcysteine carboxyl methyltransferase (ICMT-1), which finalizes protein prenylation, distinctly enriches to ER exit sites that dynamically polarize and deliver prenylated GTPases that drive protrusion formation. Collectively, these results reveal a dynamic collaboration between lipid biosynthesis and polarized lipid modification that drives invasive protrusion formation to clear BM barriers.

## Results

### Anchor cell (AC) invasive protrusive formation requires the SREBP ortholog SBP-1

The AC is a specialized uterine cell that invades through the juxtaposed uterine and vulval BMs during an ∼90-min period in the mid-L3 larval stage to initiate uterine-vulval attachment (Sherwood and Sternberg, 2003). AC differentiation and invasion occurs in synchrony with divisions of the underlying 1° fated P6.p vulval precursor cell (VPC), which allows precise staging of invasion (Fig. 1 A). The AC is specified at the late larval L2 stage, just prior to the L2/L3 molt, and during the L3 larval stage expresses pro-invasive actin regulators and matrix remodeling proteins (P6.p 1-cell stage) until the early-to-mid L3 when the AC initiates BM beaching with a small F-actin rich invadosome protrusion that penetrates the BM (P6.p 2-cell; Fig. 1 A) (Costa et al., 2023; Hagedorn et al., 2013; Kimble, 1981). After an invadosome breaches the BM, the UNC-40 (DCC) receptor localizes to the breach site and directs lysosome exocytosis to form a large protrusion that transiently increases the size of the AC by as much as ∼40% in surface area (2-4-cell stage) (Morrissey et al., 2013; Naegeli et al., 2017). Following BM clearance, the protrusion retracts, and the AC nestles between the central vulval cells (4-cell stage). The AC expresses lipid anchored proteins that are enriched at the invasive plasma membrane and then are found in the protrusion, such as the matrix degrading GPI-anchored MMP ZMP-1, the prenylated Rho GTPases CED-10 (Rac) and MIG-2 (Rac-like), and the Ras-like GTPase RAP-1 (Fig. 1, B-D; previously quantified in (Costa et al., 2023; Hagedorn et al., 2013; Kelley et al., 2019; Naegeli et al., 2017; Wang et al., 2014a). How the AC synthesizes additional lipid membranes for protrusion formation and organizes production of lipid anchored proteins is unknown.

**Figure 1.**
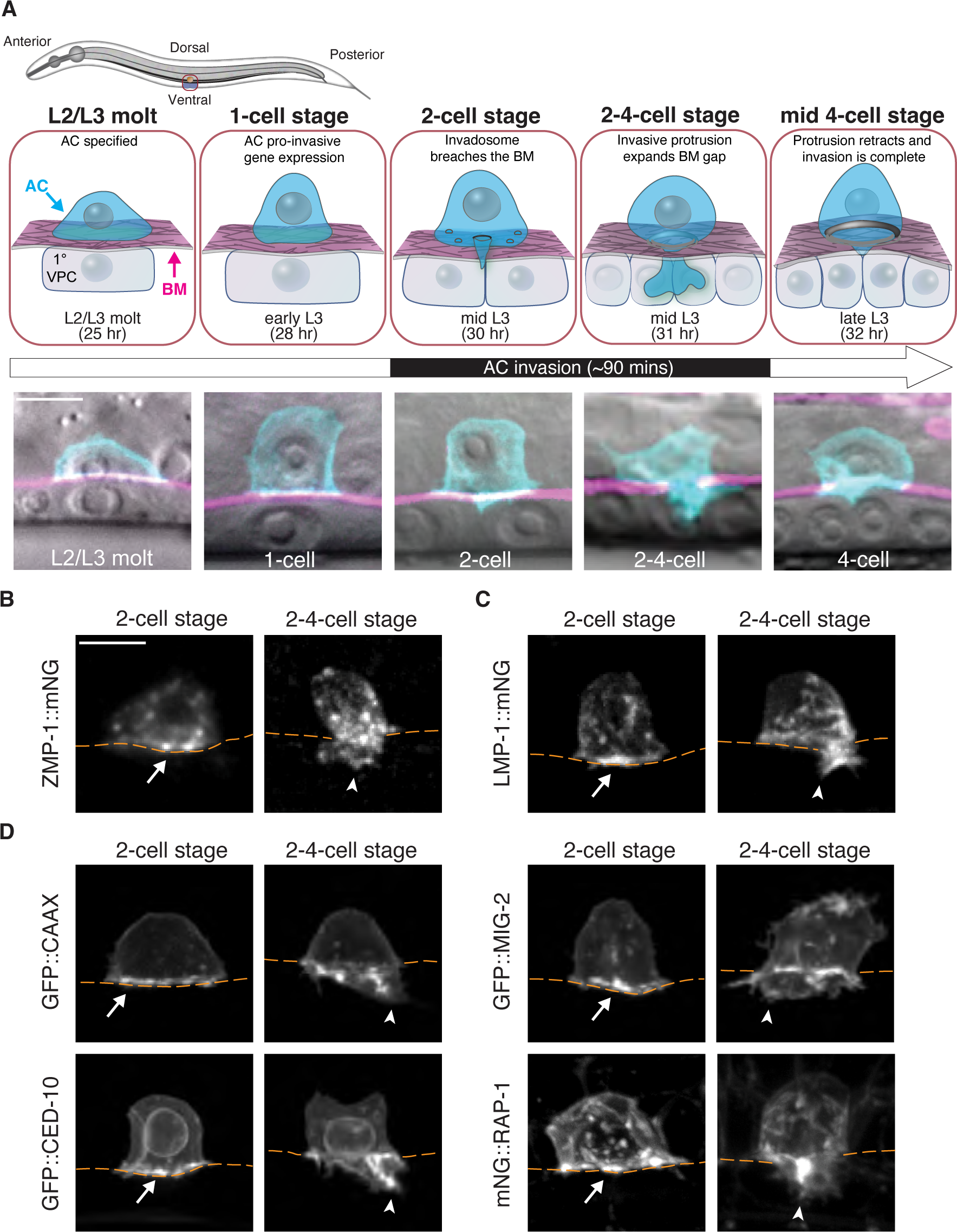
The AC invasive membrane and protrusion is enriched with lipid-modified pro-invasive proteins. **(A)** (*Top*) A schematic diagram of AC invasion. (*Bottom*) Merged differential interference microscopy (DIC) and fluorescence images (maximum intensity z-projections) showing the AC (cyan, mCherry::PLCδ^PH^), the underlying BM (magenta, LAM-1:GFP), and the vulval precursor cells (VPCs, DIC). The AC is specified at the L2/L3 molt and pro-invasive gene expression occurs at the P6.p 1-cell stage. AC invasion is initiated at the late P6.p 2-cell stage when an invadosomes breaches the BM. During the P6.p 2-4-cell stage transition, a large invasive protrusion forms at the breach site that expands the BM gap and then retracts, allowing the AC to contact the central VPCs at P6.p 4-cell stage. Timeline in h post-hatching at 20°C shown. **(B)** In the AC of wild-type animals the GPI-anchored matrix metalloproteinase ZMP-1 (ZMP-1::mNG), **(C)** lysosomes (LMP-1::mNG), **(D)** and the prenylated membrane reporter (GFP::CAAX), prenylated Rac GTPases (GFP::CED-10 and GFP::MIG-2) and Ras-like GTPase RAP-1 (mNG::RAP-1) are enriched at the AC basal plasma membrane prior to invasion (arrows; P6.p 2-cell stage) and within the AC invasive protrusion (arrowheads; P6.p 2-4-cell stage) (observed in n ≥ 10 animals for each lipid anchored protein at stage). All data shown are from two or more replicates. Scale bars, 5 µm.

Additional plasma membrane and lipid anchors could arise from external fatty acid import, stored lipids, or de novo synthesis (Martin-Perez et al., 2022). A previous investigation of carbon sources used to generate ATP for AC invasion found that the AC does not have lipid stores and does not depend on fatty acid or amino acid import (Garde et al., 2022). Instead, the AC imports glucose and uses mitochondrial oxidative phosphorylation to generate ATP that powers the invasive machinery (Garde et al., 2022). These observations suggest that the AC might generate de novo lipids by using citrate generated from the mitochondrial tricarboxylic acid cycle (TCA, also known as Krebs and citric acid cycle) to build new lipids (Martin-Perez et al., 2022).

Lipogenesis is a process by which cells convert carbohydrates to fatty acids via a series of highly regulated enzymatic reactions (see Fig. S1 A) (Ameer et al., 2014). Initially, citrate from the TCA cycle is converted to the two-carbon acetyl-CoA, which is then transformed to malonyl-CoA. Fatty acid synthesis enzymes then facilitates the production of lipids that serve as the precursor for glycerophospholipids (hereafter phospholipids) that form the bulk of membranes in the cell and specialized sphingolipids that regulate membrane stability, lipid raft formation, vesicle and protein trafficking, and signaling (Fig. S1 A)(Ameer et al., 2014; Cockcroft, 2021; Goni, 2022; Watts and Ristow, 2017). In many organisms, including *C. elegans*, the transcription of lipid biosynthetic enzymes is regulated by the SREBP family (sterol regulatory element-binding protein, *C. elegans sbp-1*) (Bertolio et al., 2019; McKay et al., 2003; Sun et al., 2020).

We hypothesized that if lipogenesis were required to build the invasive protrusion, then AC-specific loss of key lipogenesis enzymes would show defective invasion. To determine whether SBP-1 and de novo lipid biosynthesis promotes AC invasive protrusion formation, we used an AC-specific RNAi strain and targeted the *sbp-1* gene (see Table S1; methods)(Garde et al., 2022). Consistent with a possible role in invasive protrusion regulation, AC-specific reduction of *sbp-1* severely disrupted invasion (over 60% of ACs unable to remove BM by P6.p 4-cell stage; Table S2; Fig. 2 A). An *sbp-1* null mutant is lethal (McKay et al., 2003), however, the slow growing viable *sbp-1(ep79)* mutant (Liang et al., 2010) also showed invasion defects (Table S2). Examination of AC-specific membrane localized mCherry fluorophore (*cdh-3p*::mCherry::PLCδ^PH^) in the AC-specific RNAi strain, revealed that loss of *sbp-1* dramatically reduced the formation and growth of the invasive protrusion (Fig. 2 B; Fig. S2; Video 1). These results suggest that de novo lipid synthesis regulated by SBP-1 is required for invasive protrusion formation.

**Figure 2.**
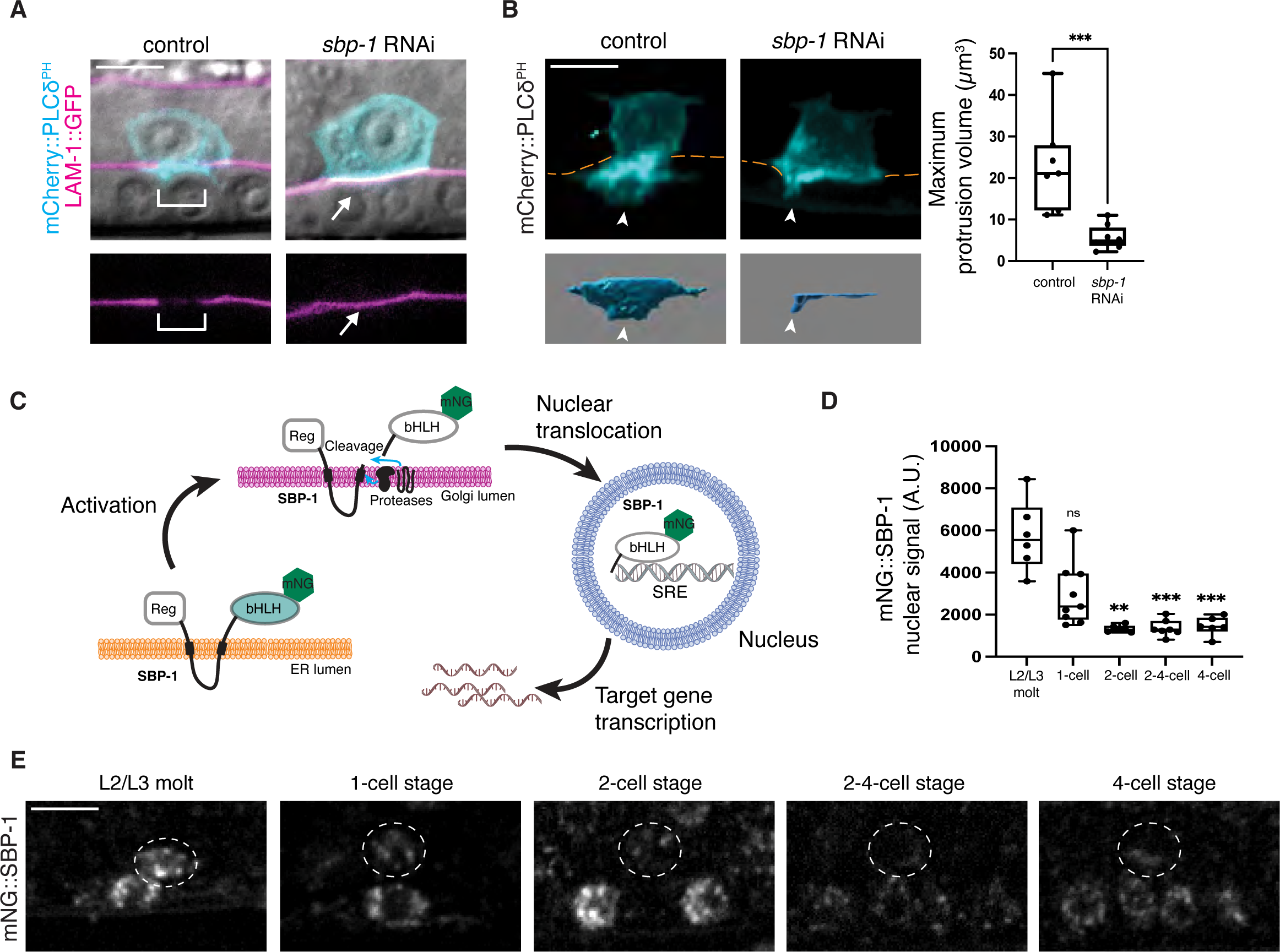
SREBP (SBP-1) promotes AC invasion and invasive protrusion formation. **(A)** (*Top*) Merged DIC and fluorescence images (maximum intensity z-projections) of the AC (cyan, mCherry::PLCδ^PH^) and the underlying BM (magenta, LAM-1::GFP) at the P6.p 4-cell stage in a control (empty RNAi vector treated) and an AC-specific *sbp-1* RNAi treated animal. (*Bottom*) Fluorescence images showing BM breach (brackets) in a control animal and lack of breach after *sbp-1* RNAi (arrows). **(B)** (*Left*) Maximum intensity z-projected fluorescence images showing the AC (cyan, mCherry::PLCδ^PH^) in a control and an AC-specific *sbp-1* RNAi treated animal (BM, orange dashed lines) with isosurfaces of the AC invasive protrusion (below, arrowheads) at the time of maximum protrusion volume (see Video 1). (*Right*) Boxplot shows the maximum invasive protrusion volume in control and *sbp-1* RNAi treated animals (n ≥ 7 animals per condition, *** p ≤ 0.001, Mann-Whitney U test). For this and all subsequent boxplots, box edges indicate the 25^th^ and 75^th^ percentiles, whiskers the maximum and minimum values, and the line within the box the median value. **(C)** A schematic diagram illustrating AC SBP-1 proteolytic activation and entry into nucleus where the SBP-1 bHLH domain binds sterol regulatory elements (SREs) to transcribe target genes. **(D)** Boxplot of mNG::SBP-1 mean fluorescence intensity in the AC nucleus from the L2/L3 molt stage to the P6.p 4-cell stage (n ≥ 6 animals per developmental stage, ** p ≤ 0.01, *** p ≤ 0.001, ns (not statistically significant), p > 0.05 One-way ANOVA followed by Kruskal-Wallis test and Dunn’s multiple comparisons test). **(E)** Sum intensity z-projected fluorescence images of mNG::SBP-1 in the AC nucleus (white dashed circle) from the L2/L3 molt (time of AC specification) to the P6.p 4-cell stage. All data are from two or more replicates. Scale bars, 5µm.

SREBP proteins reside on the ER membrane in their inactive state. However, when activated, SREBP translocates to the Golgi, where two Golgi proteases cleave SREBP and release the N-terminal DNA binding basic-helix-loop-helix (bHLH) domain, which enters the nucleus and regulates lipogenic gene expression (Fig. 2 C) (DeBose-Boyd and Ye, 2018). To visualize both the expression and activity of the *C. elegans* SBP-1 protein in the AC, we used CRISPR-Cas9 mediated genome editing to endogenously tag the *sbp-1* locus with the fluorophore mNeonGreen (mNG) at the N-terminus (mNG::SBP-1). Examination of endogenous SBP-1 in the AC revealed that SBP-1 strongly localized to the AC nucleus at the L2/L3 molt and during the early P6.p 1-cell stage, but then nuclear levels diminished rapidly after the 1-cell stage and through the time of invasion (Fig. 2 D and E). This suggests that SBP-1 is active in the AC ∼4-5 h before invasion to promote the expression of lipogenic genes required for AC invasive protrusion formation and BM breaching.

### SBP-1 regulates AC expression of key lipogenic enzymes that promote invasion

The early localization of SBP-1 in the AC nucleus suggested that SBP-1 might promote the expression of lipogenic enzymes to support invasive protrusion formation. To test this, we used AC-specific RNAi to reduce the expression of the key lipogenic transcriptional targets of SBP-1, *pod-2* (vertebrate ACC) and *fasn-1* (vertebrate FASN), which play crucial roles in the initial steps of de novo fatty acid synthesis (Fig. S1 A and Fig. 3 A) (Rohrig and Schulze, 2016; Watts and Ristow, 2017). Reduction of *pod-2* and *fasn-1* resulted in significant AC invasion defects at the P6.p 4-cell stage (∼40% and 55% respectively; Tables S1 and S2), consistent with a key role downstream of SBP-1 in invasive protrusion formation. We examined endogenously tagged GFP::POD-2 (Starich et al., 2020) and generated an endogenously tagged FASN-1::mNG. Both enzymes were highly expressed and enriched in the AC compared to neighboring non-invading uterine cells with expression increasing significantly from the P6.p 1-cell stage to the P6.p 2-4-cell stage at the time of protrusion formation (Fig. S3). RNAi mediated reduction of the SBP-1 protein also significantly reduced the levels of GFP::POD-2 and FASN-1::mNG in the AC (Fig. 3 B). These observations suggest that SBP-1 might regulate protrusion formation through transcriptional regulation of *pod-2* and *fasn-1* in the AC.

**Figure 3.**
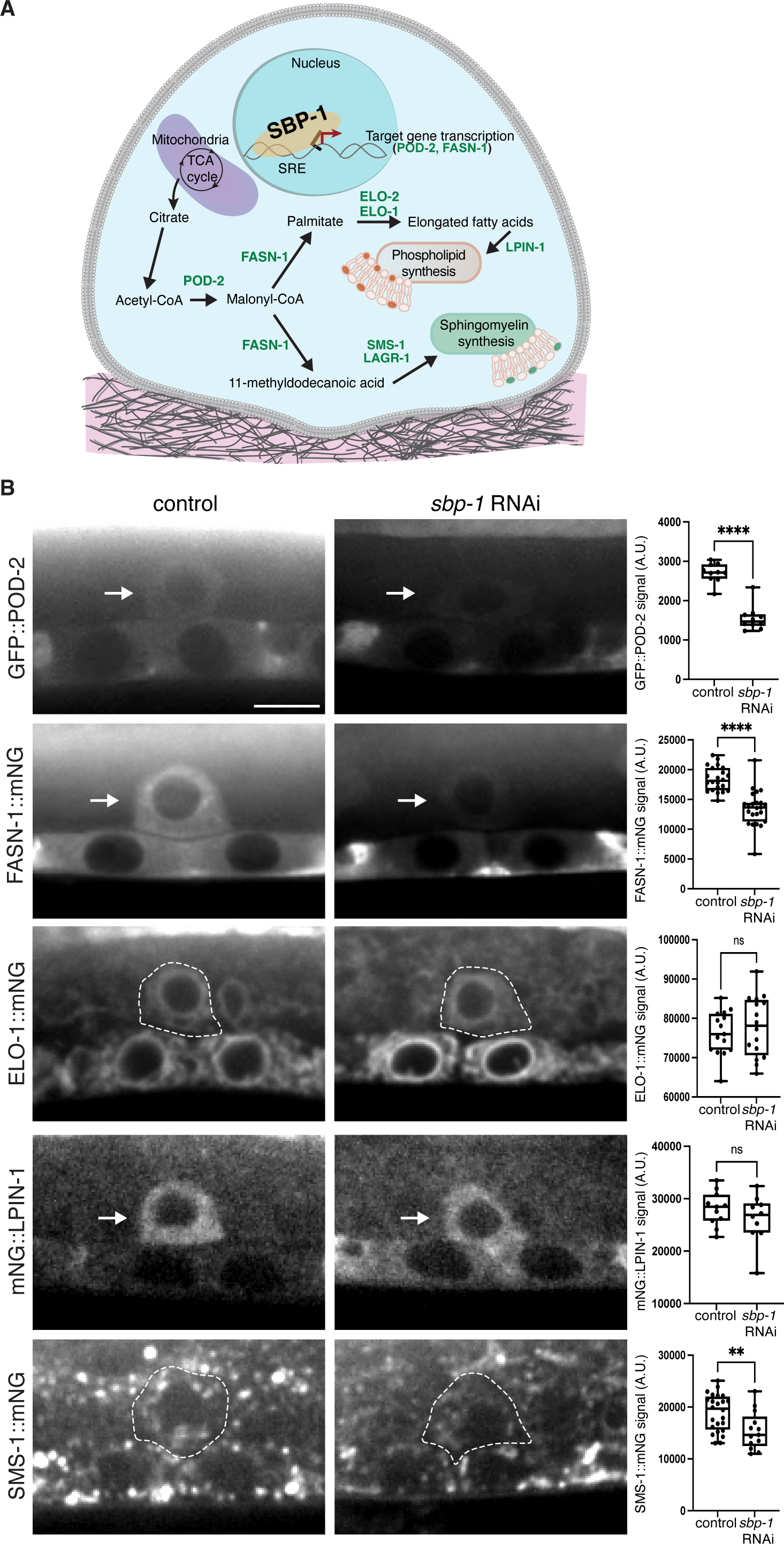
AC expression and SBP-1 regulation of *pod-2*, *fasn-1, elo-1, lpin-1, and sms-1*. **(A)** A schematic diagram showing the role of the transcription factor SBP-1 in regulating phospholipid and sphingomyelin synthesis in the AC (refer to Fig. S1 A for complete pathway and Table S2 for RNAi screen results). **(B)** Sum intensity z-projection images of POD-2 (GFP::POD-2), FASN-1 (FASN-1::mNG), ELO-1 (ELO-1::mNG), LPIN-1 (mNG::LPIN-1), and SMS-1 (SMS-1::mNG) in control and *sbp-1* RNAi animals at the P6.p 2-cell stage. POD-2, FASN-1, and LPIN-1 are found in the AC cytoplasm (arrows, ACs). ELO-1 and SMS-1 are ER and Golgi-resident proteins, respectively (ACs outlined with white dashed line). (*Right*) Boxplots shows the mean fluorescence intensity of each protein in the AC of control and *sbp-1* RNAi treated animals (n ≥ 10 animals per condition, ** p ≤ 0.01, **** p ≤ 0.0001, ns (not statistically significant), p > 0.05, unpaired two-tailed Student’s t-test and Mann-Whitney U test). All data are from two or more replicates. Scale bar, 5µm.

### De novo phospholipid and sphingolipid synthesis is required for AC invasion

Key outputs of the SBP-1 targets POD-2 and FASN-1 are palmitate, which can go on to form phospholipids and triacylglyceride for storage in lipid droplets, and 11-methyldodecanoic acid, which is a core precursor for *C. elegans* sphingolipids (Fig. 3 A and Fig. S1 A) (Watts and Ristow, 2017). As the AC does not form lipid droplets (Garde et al., 2022), it suggested that SBP-1 could promote invasive protrusion formation via phospholipid and sphingolipid production. To test this, we used AC-specific RNAi to knock down 20 key genes involved in phospholipid and sphingolipid synthesis (Table S2 and Fig. S1 A (Watts and Ristow, 2017)). RNAi depletion of two sphingolipid biosynthetic encoding enzymes, *lagr-1* and *sms-1*, significantly perturbed AC invasion (∼35% and 40% respectively) and loss of a third, *hyl-1* trended towards significance (Table S2 and Fig. 3 A). Loss of the fatty acid elongases *elo-1* and *elo-2*, as well as *lpin-1,* which synthesizes diacylglycerol for phospholipid production, also resulted in significant invasion defects (Table S2 and Fig. 3 A). We examined endogenously tagged ELO-1::mNG (Costa et al., 2023), and generated endogenous mNG knock-in strains to observe mNG::LPIN-1 and SMS-1::mNG. All enzymes were present at high levels in the AC and levels peaked near the time of protrusion formation (Fig. 3 B and Fig. S3). The enzymes were also localized to expected subcellular regions: ELO-1, the endoplasmic reticulum; LPIN-1, the cytosol; and SMS-1, in diffuse puncta consistent with dispersed Golgi stacks (Fig. 3 B) (Costa et al., 2023; D’Angelo et al., 2018; Ding et al., 2023; Naegeli et al., 2017). Notably, RNAi targeting *sbp-1* did not affect ELO-1 or LPIN-1 levels, and only modestly reduced SMS-1 (Fig. 3 B). These results are consistent with previous studies indicating that SBP-1 regulates key upstream de novo fatty acid synthesis enzymes, but not all downstream lipid synthesis genes (Nomura et al., 2010). These observations suggest that de novo produced phospholipids and sphingolipids could promote invasive protrusion formation.

### Phospholipid synthesis builds lysosome stores that construct the invasive protrusion

To investigate a role for phospholipid production in invasive protrusion formation, we knocked down *lpin-1* in the AC-specific RNAi strain (Table S1) and found that loss of *lpin-1* significantly reduced the formation of the invasive protrusion (Fig. 4 A; Fig. S2 B; Video 2). Examination of lysosomes (AC expressed LMP-1::mNG; see methods) revealed that loss of *lpin-1* reduced formation of lysosome stores prior to invasive protrusion formation (Fig. 4 B). RNAi mediated reduction of *lpin-1*, however, did not alter polarization of the prenylated Rac GTPases CED-10 and MIG-2 or the GPI-anchored ZMP-1 at the initiation of invasive protrusion formation (late P6.p 2-cell and 2-4 cell stage; Fig. 4 C and D). These results suggest that AC phospholipid production promotes invasive protrusion formation by contributing to lysosome stores that construct the protrusion but does not significantly regulate GTPase or ZMP-1 localization.

**Figure 4.**
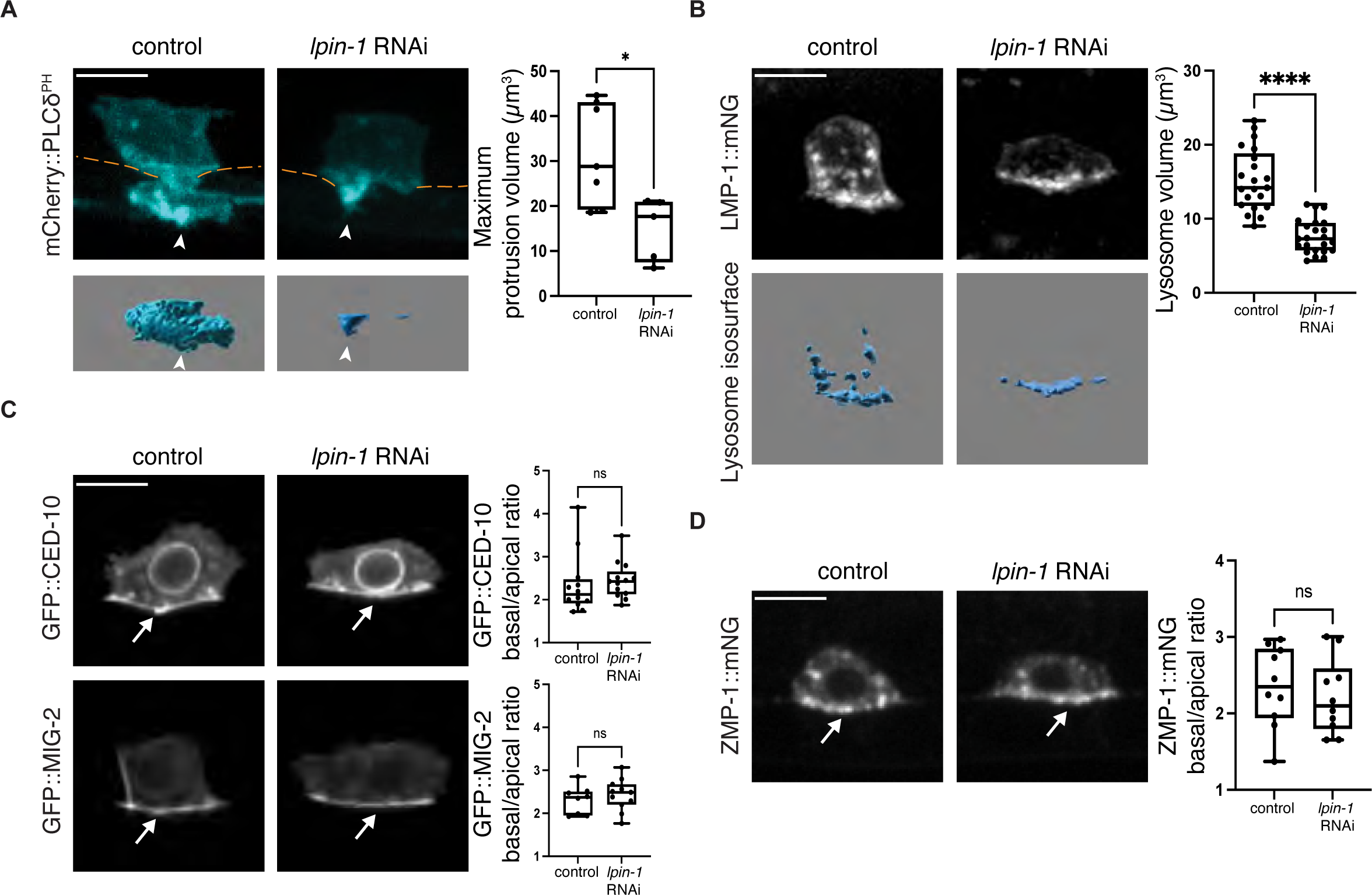
LPIN-1 promotes lysosome stores for invasive protrusion formation. **A)** (*Left*) Maximum intensity z-projected fluorescence images showing the AC (cyan, mCherry::PLCδ^PH^) in a control and an AC-specific *lpin-1* RNAi treated animal (BM, orange dashed lines) with isosurfaces of the AC invasive protrusion (below, arrowheads) at the time of maximum protrusion volume (see Video 2). (*Right*) Boxplot of the maximum invasive protrusion volume in control and *lpin-1* RNAi treated animals (n ≥ 5 animals, * p ≤ 0.05, Mann-Whitney U test). **(B)** (*Left*) Fluorescence and isosurface images showing the AC lysosomes (LMP-1::mNG) in a control and *lpin-1* RNAi treated animal at the initiation of protrusion formation (late P6.p 2-cell stage and 2-4 cell stage). (*Right*) Boxplot of AC lysosome volume in control and *lpin-1* RNAi treated animals (n ≥ 21 animals per condition, **** p ≤ 0.0001, unpaired two-tailed Student’s t-test). **(C)** (*Left)* Sum intensity z-projected images show the polarized localization of GFP::CED-10 (Rac) and GFP::MIG-2 (Rac-like) at the AC invasive plasma membrane (arrow) in control and *lpin-1* RNAi treated animals at the late P6.p 2-cell stage at the initiation of protrusion formation. (*Right*) Boxplot showing the basal/apical ratio of GFP::CED-10 and GFP::MIG-2 (n ≥ 8 animals per condition, ns (not statistically significant), p > 0.05, Mann-Whitney U test). **(D)** (*Left)* Sum intensity z-projections of ZMP-1::mNG fluorescence showing the invasive membrane enrichment (arrow) in a control and a *lpin-1* RNAi treated animal at the initiation of invasive protrusion formation (late P6.p 2-cell stage and 2-4 cell stage). *(Right)* Boxplot showing the AC basal/apical ratio of ZMP-1::mNG fluorescence intensity in control and *lpin-1* RNAi treated animals (n ≥ 10 animals per condition, ns (not statistically significant), p > 0.05, unpaired two-tailed Student’s t-test). All data are from two or more replicates. Scale bars, 5µm.

### Sphingomyelin has multiple functions in building the AC’s invasive protrusion

The sphingomyelin synthase SMS-1 generates sphingomyelin and was the most downstream enzyme in sphingolipid synthesis whose loss perturbed AC invasion (Table S2 and Fig. S1 A) (Watts and Ristow, 2017). This suggested that sphingomyelin might be crucial for invasive protrusion formation. Sphingomyelin composes ∼ 5% of membrane lipids and is found in the plasma membrane and endolysosome, where it plays roles in signaling, vesicle and protein trafficking, membrane stability, and lysosome function (Duran et al., 2012; Goni, 2022; Tang et al., 2022). We knocked down *sms-1* in the AC-specific RNAi strain and found that loss of *sms-1* diminished the formation and growth of the invasive protrusion (Fig. 5 A; Fig. S2 B; Video 3). Notably, loss of *sms-1* reduced lysosome stores that build the protrusion (Fig. 5 B). Rac GTPase enrichment at the initiation of protrusion formation, however, was unaltered (Fig. 5 C). Sphingomyelin is a key component of lipid rafts (Bieberich, 2018), which partition GPI-anchored proteins and the vertebrate UNC-40 receptor ortholog DCC (Herincs et al., 2005; Hernaiz-Llorens et al., 2021; Sangiorgio et al., 2004). We found that loss of *sms-1* reduced the enrichment of the GPI-anchored ZMP-1 protein at the site of invasive protrusion formation (Fig. 5 D). Further, ventral view time-lapse analysis along the BM plane revealed that *sms-1* loss also reduced UNC-40::GFP receptor concentration at the BM breach site (Fig. 5 E), where UNC-40 directs lysosome exocytosis to form the invasive protrusion (Hagedorn et al., 2013; Morrissey et al., 2013; Naegeli et al., 2017). These results suggest that sphingomyelin is crucial for forming lysosomes that build the protrusion, for regulating ZMP-1 localization for the BM degradation, and for promoting UNC-40 localization that directs protrusion formation. Together, these studies build a network of de novo lipid synthesis genes acting downstream of SBP-1 that mediate invasive protrusion formation and function (Fig. 5 F).

**Figure 5.**
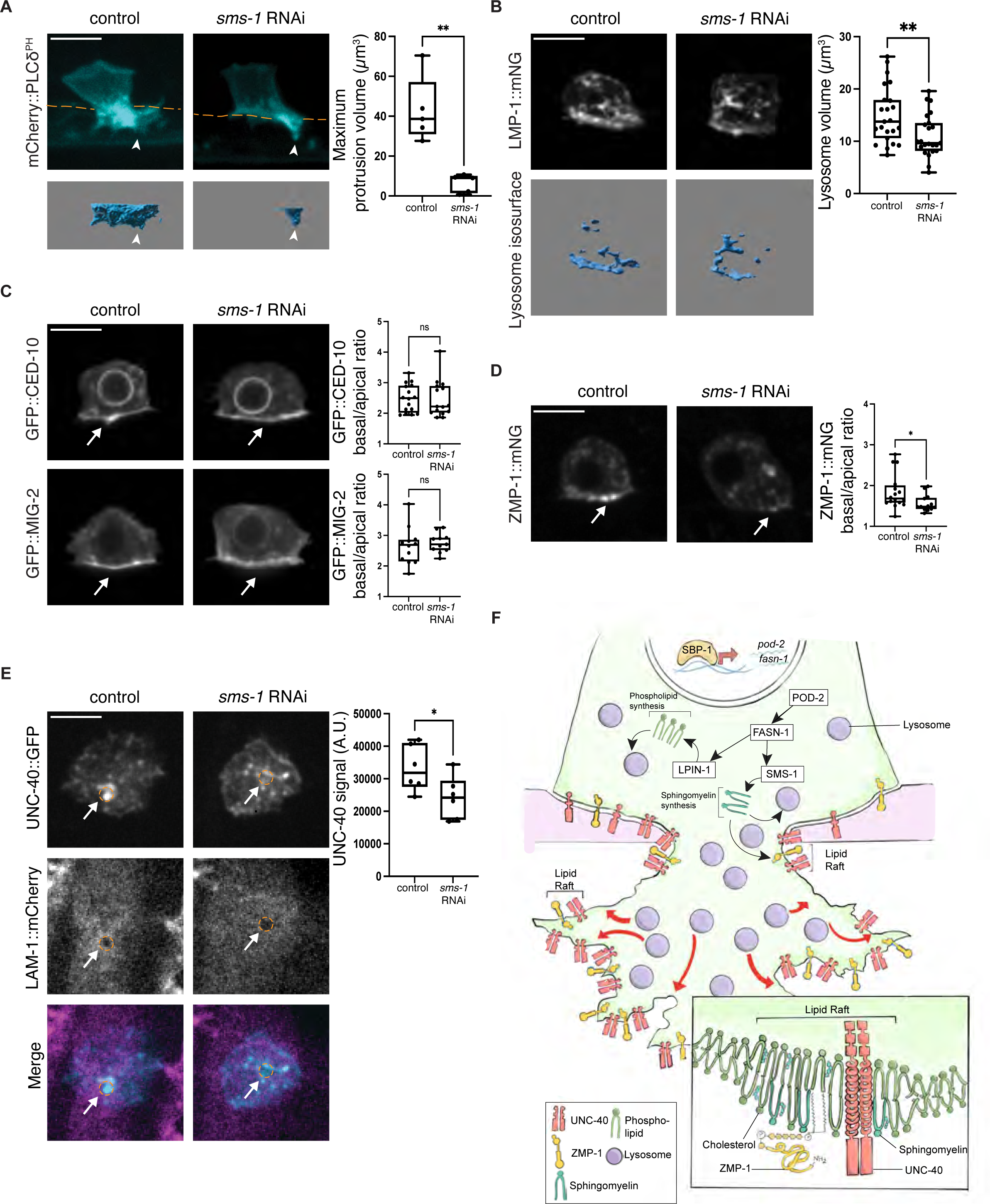
SMS-1 promotes formation of lysosome stores and localization of UNC-40 and ZMP-1. **(A)** (*Left*) Maximum intensity z-projected fluorescence images showing the AC (cyan, mCherry::PLCδ^PH^) in a control and an AC-specific *sms-1* RNAi treated animal (BM, orange dashed lines) with isosurfaces of the AC invasive protrusion (below, arrowheads) at the time of maximum protrusion volume (see Video 3). (*Right*) Boxplot of the maximum invasive protrusion volume in control and *sms-1* RNAi treated animals (n ≥ 5 animals per condition, ** p ≤ 0.01, Mann-Whitney U test). **(B)** (*Left*) Sum intensity z-projected fluorescence images and isosurfaces showing the AC lysosomes (LMP-1::mNG) in a control and *sms-1* RNAi treated animal at the initiation of protrusion formation. (*Right*) Boxplot of the AC lysosome volume in control and *sms-1* RNAi treated animals (n ≥ 23 animals per condition, ** p ≤ 0.01, unpaired two-tailed Student’s t-test). **(C)** (*Left*) Sum intensity z-projected fluorescence images of GFP::CED-10 and GFP::MIG-2. Arrows show enrichment at the AC invasive membrane in a control and *sms-1* RNAi treated animal at the time of invasive protrusion initiation (late P6.p 2-cell and 2-4 cell). (*Right*) Boxplots show the basal/apical ratio of GFP::CED-10 and GFP::MIG-2 fluorescence intensity in control and *sms-1* RNAi treated animals (n ≥ 11 animals per condition, ns (not statistically significant), p > 0.05, Mann-Whitney U test). **(D)** (*Left*) Sum intensity fluorescence images of the GPI-anchored matrix metalloproteinase ZMP-1::mNG showing the AC invasive membrane (arrows) in a control and a *sms-1* RNAi treated animal at the initiation of invasive protrusion formation. (*Right*) Boxplot showing the AC basal/apical ratio of ZMP-1::mNG fluorescence intensity in control and *zmp-1* RNAi treated animals (n ≥ 13 animals per condition, * p ≤ 0.05, unpaired two-tailed Student’s t-test). **(E)** (*Left*) Ventral view of the AC-BM interface (BM visualized with LAM-1::mCherry) showing UNC-40::GFP enrichment (arrows) at the initial BM breach (orange dotted lines) in a control and a *sms-1* treated RNAi animal. (*Right*) Boxplot shows UNC-40::GFP mean fluorescence intensity at the initial BM breach (n= 6 animals per condition, * p ≤ 0.05, Mann-Whitney U test). **(F)** A schematic diagram summarizing the roles of the transcription factor SBP-1, the fatty acid synthesis enzymes POD-2 and FASN-1, and phospholipid synthesizing LPIN-1 and sphingomyelin catalyzing SMS-1 in AC invasive protrusion formation. All data in figure are from two or more replicates. Scale bars, 5µm.

### The mevalonate pathway promotes lysosome stores and prenylated protein localization

In addition to being converted into fatty acids, acetyl-CoA can also be used to generate products of the mevalonate pathway. The mevalonate pathway converts acetyl-CoA into five-carbon branched isoprene groups that are turned into sterols and prenol lipids, such as cholesterol, coenzyme Q, dolichol, and prenylated anchors (15-carbon farnesyl and 20-carbon geranylgeranyl) for proteins such as small GTPases (Fig. 6 A and Fig. S1 B) (Rauthan and Pilon, 2011). In mammals, the mevalonate pathway produces cholesterol and SREBP is a sensor of cholesterol levels and transcriptionally regulates the rate limiting enzyme of the mevalonate pathway, HMG-CoA reductase (HMGCR, *C. elegans hmgr-1)*(Rauthan and Pilon, 2011). C*. elegans* do not synthesize cholesterol (Vinci et al., 2008; Watts and Ristow, 2017) and instead acquire cholesterol from feeding, and it is unknown if SBP-1 regulates *hmgr-1* expression (Rauthan and Pilon, 2011).

**Figure 6.**
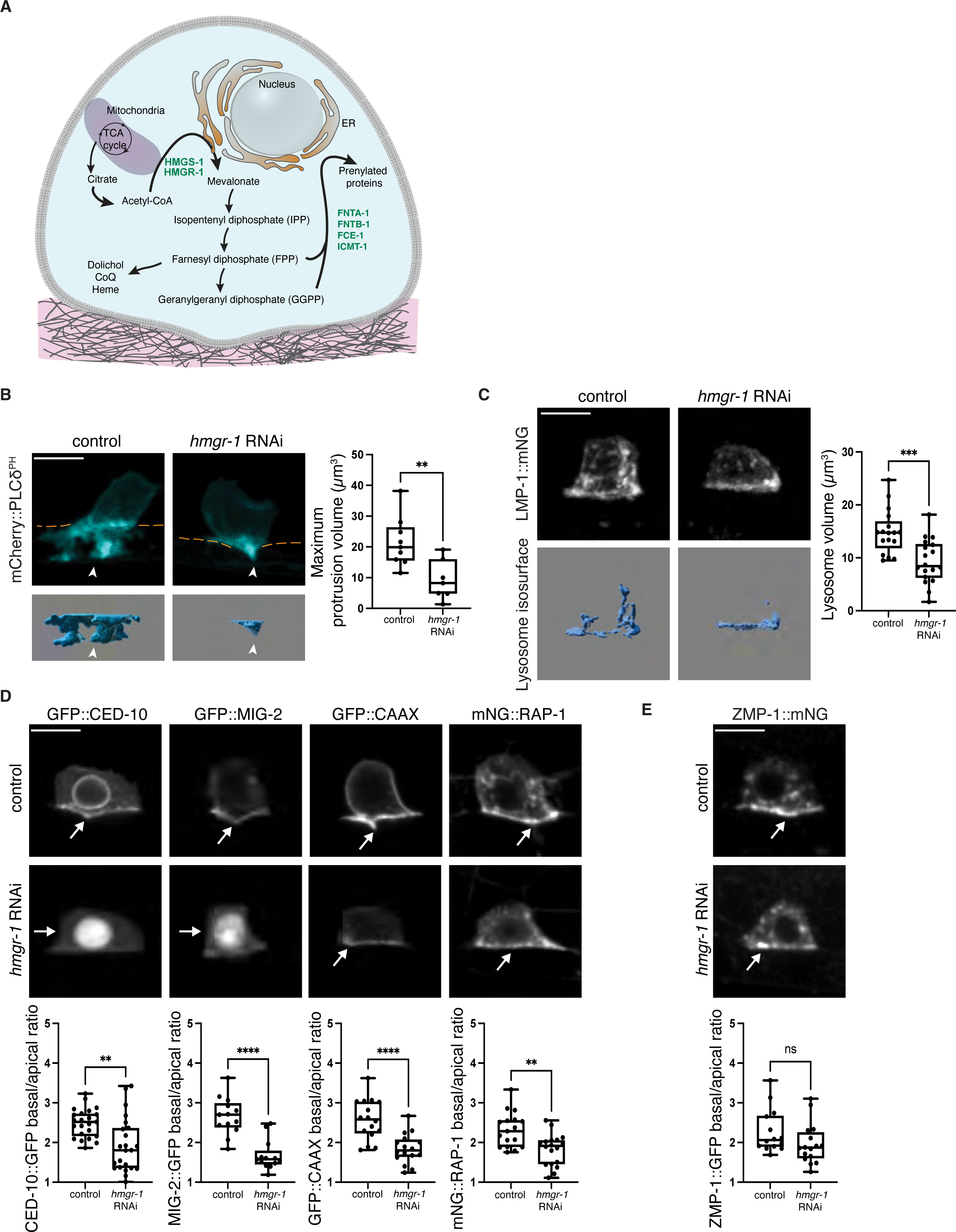
HMGR-1 promotes lysosome stores, GTPase accumulation and invasive protrusion formation. **(A)** A schematic diagram showing the mevalonate pathway in the AC (refer to Fig. S1 B for details on the pathway and Table S2 for RNAi screen results). **(B)** (*Left*) Maximum intensity z-projected fluorescence images showing the AC (cyan, mCherry::PLCδ^PH^) in a control and an AC-specific *hmgr-1* RNAi treated animal (BM, orange dashed lines) with isosurfaces of the AC invasive protrusion (below, arrowheads) at the time of maximum protrusion volume (see Video 4). (*Right*) Boxplot of the maximum invasive protrusion volume in control and *hmgr-1* RNAi treated animals (n ≥ 7 animals per condition, ** p ≤ 0.01, Mann-Whitney U test). **(C)** (*Left*) Sum intensity z-projected fluorescence images and isosurfaces showing the AC lysosomes (LMP-1::mNG) in a control and an *hmgr-1* RNAi treated animal at the initiation of protrusion formation. (*Right)* Boxplot of the AC lysosome volume of control and *hmgr-1* RNAi animals (n ≥ 17 animals per condition, *** p ≤ 0.001, unpaired two-tailed Student’s t-test) **(D)** (*Top*) Sum intensity z-projected images showing invasive polarization (arrows) of GFP::CED-10, GFP::MIG-2, GFP::CAAX, and mNG::RAP-1 at the AC plasma membrane in control and *hmgr-1* RNAi animals at initiation of invasive protrusion formation. (*Bottom*) Boxplots showing the AC basal/apical ratio of each protein’s fluorescence intensity in control and after *hmgr-1* RNAi treatment (n ≥ 13 animals per condition, ** p ≤ 0.01, **** p ≤ 0.0001, unpaired two-tailed Student’s t-test). **(E)** (*Top*) Sum intensity projection of ZMP-1::mNG shows ZMP-1 enrichment at the AC basal invasive plasma membrane (arrows) in a control and an *hmgr-1* RNAi treated animal at the initiation of invasive protrusion formation. (*Bottom*) Boxplot showing the AC basal/apical ratio of ZMP-1::mNG fluorescence intensity in control and *hmgr-1* RNAi treated animals (n ≥ 17 animals per condition, ns (not statistically significant), p > 0.05, unpaired two-tailed Student’s t-test. All data are from two or more replicates. Scale bars, 5µm.

The rate limiting enzyme of the mevalonate pathway is HMG-CoA reductase (*C. elegans hmgr-1*). HMG-CoA synthase (*hmgs-1*) is also a key upstream regulator (Fig. S1 B) (Watts and Ristow, 2017). Using the AC-specific RNAi strain, we found that loss of *hmgr-1* and *hmgs-1* both strongly blocked AC invasion (Table S2). Time-lapse analysis of protrusion formation after RNAi targeting *hmgr-1* revealed a significant reduction in protrusion formation and growth rate (Fig. 6 B; Fig. S2 B; Video 4). Examination of key regulators of protrusion formation and function after *hmgr-1* loss further revealed that HMGR-1 is required for generating lysosome stores prior to invasion (Fig. 6 C), for the polarized enrichment of the prenylated small GTPases CED-10, MIG-2, and RAP-1, and GFP::CAAX (Fig. 6 D), but not for the enrichment of the GPI-anchored ZMP-1 (Fig. 6 E). Together these results implicate a crucial role for HMGR-1/the mevalonate pathway in targeting prenylated GTPases to the invasive protrusion, as well as the generation of lysosomes that form the protrusion.

We were next interested in understanding what lipid derivatives that are formed from the product of HMGR-1 promote lysosome construction and GTPase localization. We thus used the AC-specific RNAi strain and knocked down 10 key enzymes acting downstream of HMGR-1 that construct diverse lipid products (Table S2 and Fig. S1 B) (Rauthan and Pilon, 2011). Notably, loss of polyprenol reductase (*B0024.13*), which catalyzes the conversion of polyprenol to dolichol, strongly perturbed AC invasion. Prenylation (farnesyl and geranylgeranyl) was also strongly implicated, as RNAi mediated loss of the prenyltransferases *fnta-1* and *fntb-*1 and the isoprenylcysteine carboxylmethyltransferase *icmt-1*, which catalyzes the last step of prenylation, significantly disrupted AC invasion (Fig. 6 A; Fig. S1 B; and Table S2) (Wang and Casey, 2016). We thus further examined the polyprenol reductase *B0024.13* and *icmt-1* (the final step in prenylation) and found that loss of *B0024.13* reduced lysosome stores, while RNAi targeting of *icmt-1* decreased the invasive membrane localization of the prenylated Rac CED-10 (Fig. S4, A-C). These results suggest that dolichol, which plays a key role in N-linked glycosylation, is required for forming lysosome stores that construct the protrusion, while prenylation is crucial for prenylated GTPase localization for protrusion formation.

### ER-localized HMGR-1 and protein prenylation enzymes polarize to the AC’s invasive front

Given the unique role of HMGR-1 in regulating the localization of Rac and Ras-related GTPases in the AC, we were next interested in determining the expression and regulation of HMGR-1. HMG-CoA reductases are conserved multi-pass transmembrane ER-resident proteins (Fig. 7 A) (Schumacher and DeBose-Boyd, 2021). Using genome editing, we knocked-in mNG at the N-terminus of the *hmgr-1* locus (mNG::HMGR-1) and found that HMGR-1 localized to the peri-nuclear region in many cell types consistent with ER localization. Notably, HMGR-1 was dramatically enriched at the AC invasive side prior to and during invasion—an enrichment not observed in neighboring uterine cells (Fig. 7 B and Fig. S4 D). The mevalonate pathway in *C. elegans* does not contain the cholesterol synthesis branch and it is unknown if SBP-1 regulates *hmgr-*1/HMG-CoA reductase expression as SREBP proteins do in organisms that produce cholesterol. RNAi mediated reduction of *sbp-1* did not alter the levels of mNG::HMGR-1 in the AC (Fig. S4 E), indicating independence of HMGR-1 expression from SBP-1 regulation.

**Figure 7.**
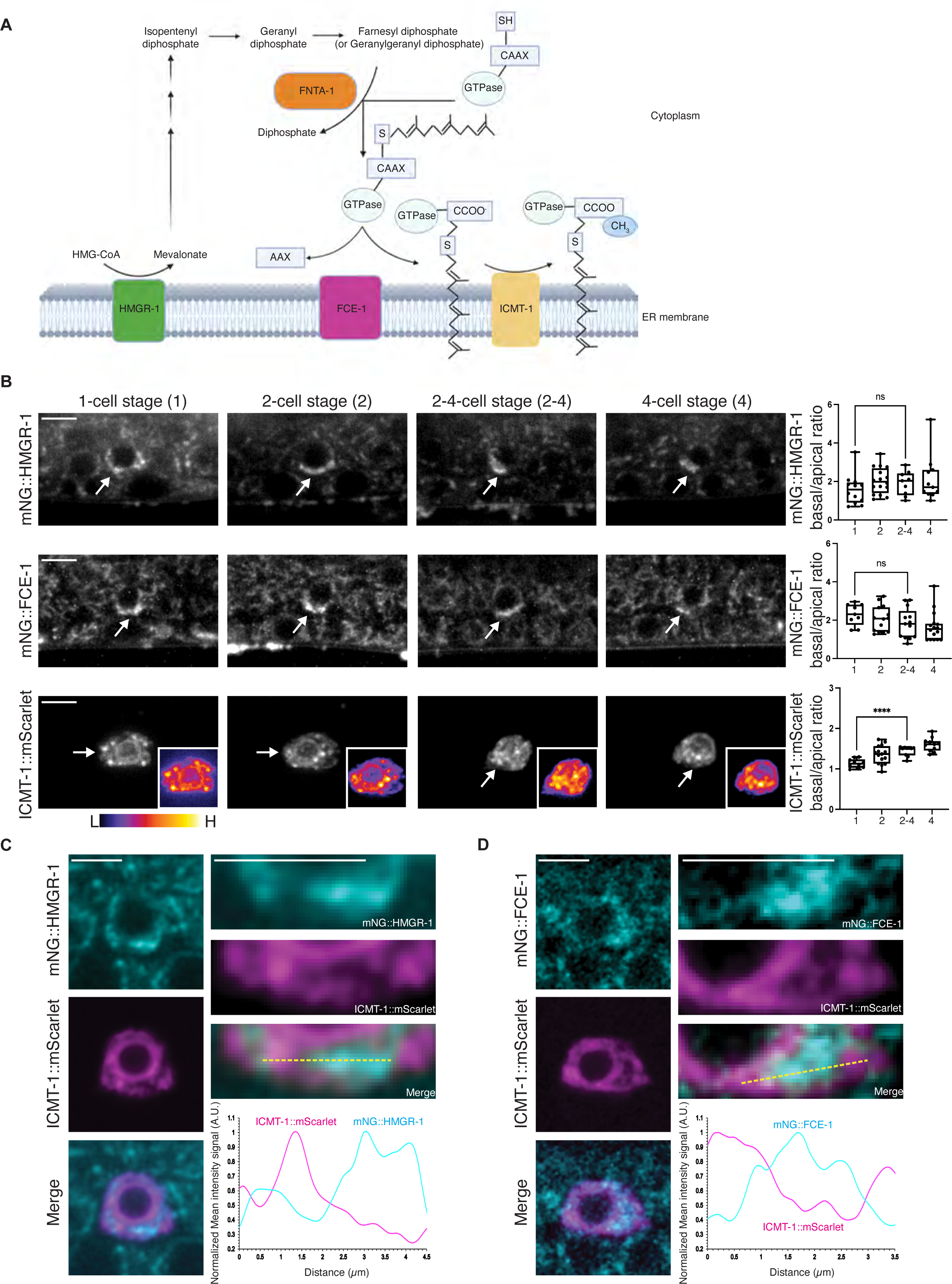
Prenylation enzymes HMGR-1, FCE-1, and ICMT-1 polarize to distinct ER subdomains. **(A)** A schematic diagram showing the mevalonate/HMGR-1 prenylation pathway (see text for detailed description). **(B)** (*Left*) Sum intensity z-projected images show AC localization and invasive enrichment (arrows) of the endogenously tagged ER-associated prenylation enzymes mNG::HMGR-1 and mNG::FCE-1, and AC expressed ICMT-1::mScarlet (*lin-29p::icmt-1::mScarlet,* inset shows spectral fluorescence-intensity map displaying the minimum and maximum pixel value range of the acquired data) from the P6.p 1-cell to 4-cell stage. (*Right*) Boxplot shows the AC basal/apical ratios of mNG::HMGR-1, mNG::FCE-1, and ICMT-1::mScarlet fluorescence intensity (n ≥ 9 animals per condition, **** p ≤ 0.0001, ns (not statistically significant), p > 0.05, One-way ANOVA followed by Brown Forsythe and Welch ANOVA tests and One-way ANOVA followed by Kruskal-Wallis test and Dunn’s multiple comparisons test). **(C)** (*Left*) Single slice confocal images of mNG::HMGR-1 (cyan), ICMT-1::mScarlet (magenta) and overlay in the AC. (*Right*) Insets highlighting the AC basal ER region. Fluorescence intensity distribution of mNG::HMGR-1 and ICMT-1::mScarlet along the yellow dashed line shows localization in different ER domains (n = 10 animals). **(D)** (*Left*) Single slice confocal images of mNG::FCE-1 (cyan), ICMT-1::mScarlet (magenta), and overlay in the AC. (*Right*) Insets highlighting the AC basal ER region. Fluorescence intensity distribution of mNG::FCE-1 and ICMT-1::mScarlet along the yellow dashed line shows localization in different ER domains (n = 10 animals). All data are from two or more replicates. Scale bars, 5µm.

UNC-6 (vertebrate netrin) is secreted by the vulval P6.p cell and its descendants and orients Rho GTPases, other actin regulators, and metabolic components of the invasive machinery to orient the invasive protrusion towards the BM (Garde et al., 2022; Hagedorn et al., 2013; Naegeli et al., 2017; Wang et al., 2014b). We thus hypothesized that HMGR-1 might also be a component of the invasive machinery polarized by UNC-6. Consistent with this notion, HMGR-1 was no longer strongly enriched towards the BM in *unc-6(ev400)* null mutant animals (Fig. S5 A). These results suggest that HMGR-1 is a component of the invasive machinery for BM breaching.

After the action of the cytosolic prenyltransferases such as FNTA-1, prenylated proteins are processed by the ER multi-pass transmembrane resident RAS-converting CAAX endopeptidase 1 (RCE1, *C. elegans* FCE-1), which removes the-AAX residue, followed by the action of the ER-resident ICMT, which adds a methyl group onto the carboxyl terminus that increases membrane interactions of prenylated proteins (Fig. 7 A) (Wang and Casey, 2016).

Given the polarized enrichment of the ER-resident HMGR-1 protein towards the site of protrusion formation, we wanted to determine the localization of FNTA-1 and the ER-resident FCE-1 and ICMT-1 proteins. We used genome editing to tag the *fnta-1* locus at the C-terminus (FNTA-1::mNG), the *fce-1* locus at the N-terminus (mNG::FCE-1), but were unable to tag *icmt-1* at its endogenous locus and thus drove its expression under an AC-specific promoter (*lin-29p::icmt-1::mScarlet*). As expected FNTA-1::mNG was cytosolic, but levels were elevated in the AC and there was a consistent enrichment at the invasive front during BM breaching (Fig. S5 B). Like HMGR-1, both ER-resident FCE-1 and ICMT-1 were also enriched towards the AC’s invasive side (Fig. 7 B) and like HMGR-1, ICMT-1 was polarized towards the BM by UNC-6 (Fig. S5 C). Interestingly, ICMT-1 polarization was highest at the time of protrusion formation, whereas HMGR-1 and FCE-1 showed similar polarization prior to and during invasion (Fig. 7 B). In addition, co-localization of ICMT-1 with FCE-1 and HMGR-1 revealed that ICMT-1 was in a distinct domain of the ER (Fig. 7 C and D). Together, these observations indicate that the ER-localized prenylation enzymes HMGR-1 and ICMT-1 are polarized to the site of invasive protrusion formation by UNC-6 (netrin), and that the final step of prenylation mediated by ICMT-1 occurs in a distinct ER domain that becomes most polarized during protrusion formation.

### ICMT-1 concentrates at ER exit sites, which polarize towards the invasive protrusion

As ICMT-1 mediates the last step of prenylation and was polarized towards the BM within a distinct ER domain, we were next interested in determining the identity of this region. We noted that ICMT-1 concentrated in puncta throughout the ER (Fig. 8 A) and hypothesized that these might be ER exit sites to allow the rapid trafficking of fully prenylated proteins to the invasive protrusion. To test this idea, we knocked in mNG into the N-terminus of the *sec-16a.1* locus (mNG::SEC-16A.1), whose protein marks ER exit sites (Watson et al., 2006), and found an ∼100% overlap of mScarlet::ICMT-1 with mNG::SEC-16A.1 (Fig. 8 A). Consistent with a role for trafficking prenylated proteins from the ER exit sites to the cell surface, RNAi reduction of *sar-1*, a GTPase that mediates COPII vesicle formation at ER exit sites for trafficking of ER cargo to the Golgi and then cell surface (Van der Verren and Zanetti, 2023), led to retention of membrane vesicles harboring GFP::CED-10 near the invasive plasma membrane (Fig. 8 B).

**Figure 8.**
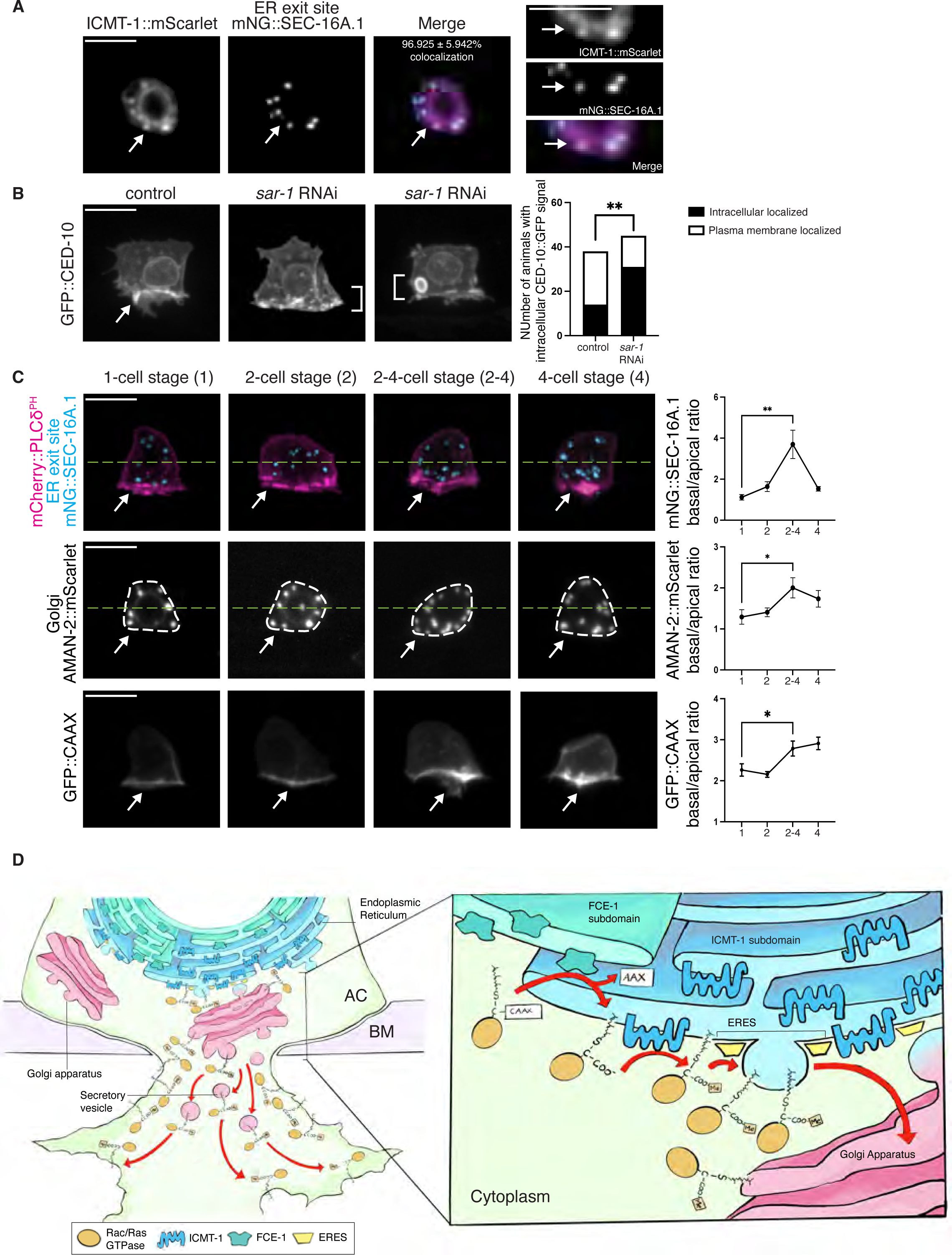
ICMT-1 localizes to ER exit sites that dynamically polarize toward the invasive protrusion. **(A)** (*Left*) Sum intensity z-projected images of ICMT-1::mScarlet (magenta), ER exit marker mNG::SEC-16A.1 (cyan), and overlay in the AC reveal co-localization at ER exit sites. The average percentage ± SD of colocalization of ER exit sites with ICMT-1 puncta is indicated (n = 17 animals). (*Right)* Insets highlight ICMT-1::mScarlet and mNG::SEC-16A.1 colocalization in the ER at the AC invasive front. **(B)** Max intensity z-projected fluorescence images of GFP::CED-10 in a control and a *sar-1* RNAi treated animal shows GFP::CED-10 localized to the invasive plasma membrane (arrow) in control animals, but found strongly in intracellular localized vesicles in *sar-1* RNAi animals (brackets). (*Right*) The relative proportion of animals with CED-10::GFP that localized predominantly to the AC basal plasma membrane or was found predominantly in the cytosol in control and *sar-1* RNAi treated animals (n ≥ 9 per condition, ** p ≤ 0.01, Fisher’s exact test). **(C)** Sum intensity z-projection showing AC ER exit sites (mNG::SEC-16A.1, Cyan; AC outlined with the plasma membrane marker mCherry::PLCδ^PH^, red), Golgi (AMAN-2::mScarlet), and prenylated GFP::CAAX in the AC from the P6.p 1-cell to 4-cell stage. Arrows indicate basal (invasive front) enrichment of each protein. (*Right)* Quantifications of the relative distribution of the ER exit site, Golgi, and plasma membrane markers in the AC from the P6.p 1-cell to 4-cell stage. For mNG::SEC-16A.1 and AMAN-2::mScarlet, the total number of puncta in the AC apical and basal halves (green dashed line) was expressed as a basal/apical ratio. The mean fluorescence intensity of GFP::CAAX was expressed as a basal/apical ratio (n ≥ 10 per condition, * p ≤ 0.05, ** p ≤ 0.01, Mann-Whitney U test, and One-way ANOVA followed by Brown Forsythe and Welch ANOVA tests). **(D)** A schematic diagram illustrating the unique ER subdomains where GTPase prenylation occurs, with ICMT-1, the final prenylation enzyme, localized to ER exit sites (ERES) that deliver prenylated proteins (*i.e,* Rac and Ras-related) to the Golgi apparatus for targeting to the invasive protrusion. All data are from two or more replicates. Scale bars, 5µm.

We next wanted to determine the localization ER exit sites and the Golgi (which has dispersed cisternae in *C. elegans*) (Sato et al., 2014), prior to and during invasion to examine the dynamics of the vesicle trafficking route during protrusion formation. Strikingly, both ER exit sites and the Golgi cisternae polarized most strongly towards the BM specifically during invasive protrusion formation (P6.p 2-4 cell stage) (Fig. 8 C). The polarization of the vesicular apparatus also coincided with the increased polarity of GTPases and GFP::CAAX (Fig. 8 C and Fig. S5 D). Taken together, these observations indicate that the ER-localized prenylation machinery within the AC is regionally organized, and that the final step of prenylation occurs at ER exit sites, which dynamically polarize with the Golgi to direct prenylated proteins to the invasive protrusion during BM breaching (Fig. 8 D).

## Discussion

Cell invasion through BM barriers is crucial for many developmental processes and immune cell trafficking and the cell invasion program is hijacked in cancer metastasis (Kelley et al., 2014; Paterson and Courtneidge, 2018). To transmigrate BMs, cells use large membrane protrusions that harbor signaling molecules and lipid anchored proteins to clear the dense, covalently cross-linked BM protein network (Naegeli et al., 2017; Nazari et al., 2023; Schoumacher et al., 2010). Although many lipid synthesis enzymes are strongly associated with cancer progress (Martin-Perez et al., 2022), due to the complexity of lipid metabolism and difficulty of studying BM invasion, it has been challenging to establish the significance, regulation, and potential roles of lipid synthesis and lipid modification within invasive cells.

Using the model system of *C. elegans* AC invasion, endogenous gene tagging, AC-specific RNAi, genetic analysis, and live-cell imaging, we show that de novo lipid synthesis is crucial for BM invasion. We discovered that the *C. elegans* SREBP transcription factor ortholog SBP-1 is transiently present at high levels in the AC nucleus ∼5 h prior to invasion and is required for invasive protrusion formation. SBP-1 promotes the expression of acetyl-CoA carboxylase ACC (*C. elegans* POD-2), the key rate limiting enzyme of de novo fatty acid synthesis, as well as the fatty acid synthase FASN-1 (Rohrig and Schulze, 2016). SREBP functions have primarily been studied in physiological and pathophysiological settings (Shimano and Sato, 2017). These findings add to a limited number of developmental studies indicating key cell-autonomous specific roles for SREBP proteins in morphogenetic cell behaviors, such as in the lipid intensive process of mouse cortical neurite plasma membrane outgrowth and *Drosophila* sensory neuron dendrite membrane expansion (Hasegawa et al., 2012; Meltzer et al., 2017; Ziegler et al., 2017). Our studies support a model where POD-2 and FASN-1 generate fatty acids within the AC that the phosphatidic acid phosphatase lipin (LPIN-1) and sphingomyelin synthase SMS-1 use to generate phospholipids and sphingomyelin that form lysosome stores that are exocytosed to generate the invasive protrusion. Further, SMS-1 also regulates the localization of the lipid raft associated DCC (UNC-40) receptor, which directs invasive protrusion formation, and the GPI-anchored ZMP-1 protease (Naegeli et al., 2017). In addition, our studies reveal that the HMG-CoA reductase HMGCR (HMGR-1), the rate limiting enzyme in the mevalonate pathway (Rauthan and Pilon, 2011), plays a crucial role in invasive protrusion formation by mediating the polarized prenylation and delivery of proteins, such as Rac GTPases to the invasive protrusion. Notably, in vertebrates the mevalonate pathway produces cholesterol, and HMGCR expression is under the control of sterol responsive SREBP transcription (DeBose-Boyd and Ye, 2018). However, *C. elegans* lacks the cholesterol-synthetic branch of the mevalonate pathway (Rauthan and Pilon, 2011), and we found that the SBP-1 does not regulate HMGR-1 expression. Together, these studies establish that an invasive cell uses a robust de novo lipid synthesis network that acts in concert with polarized prenylation to rapidly form a dynamic, specialized protrusion that removes BM barriers (Fig. 9).

**Figure 9.**
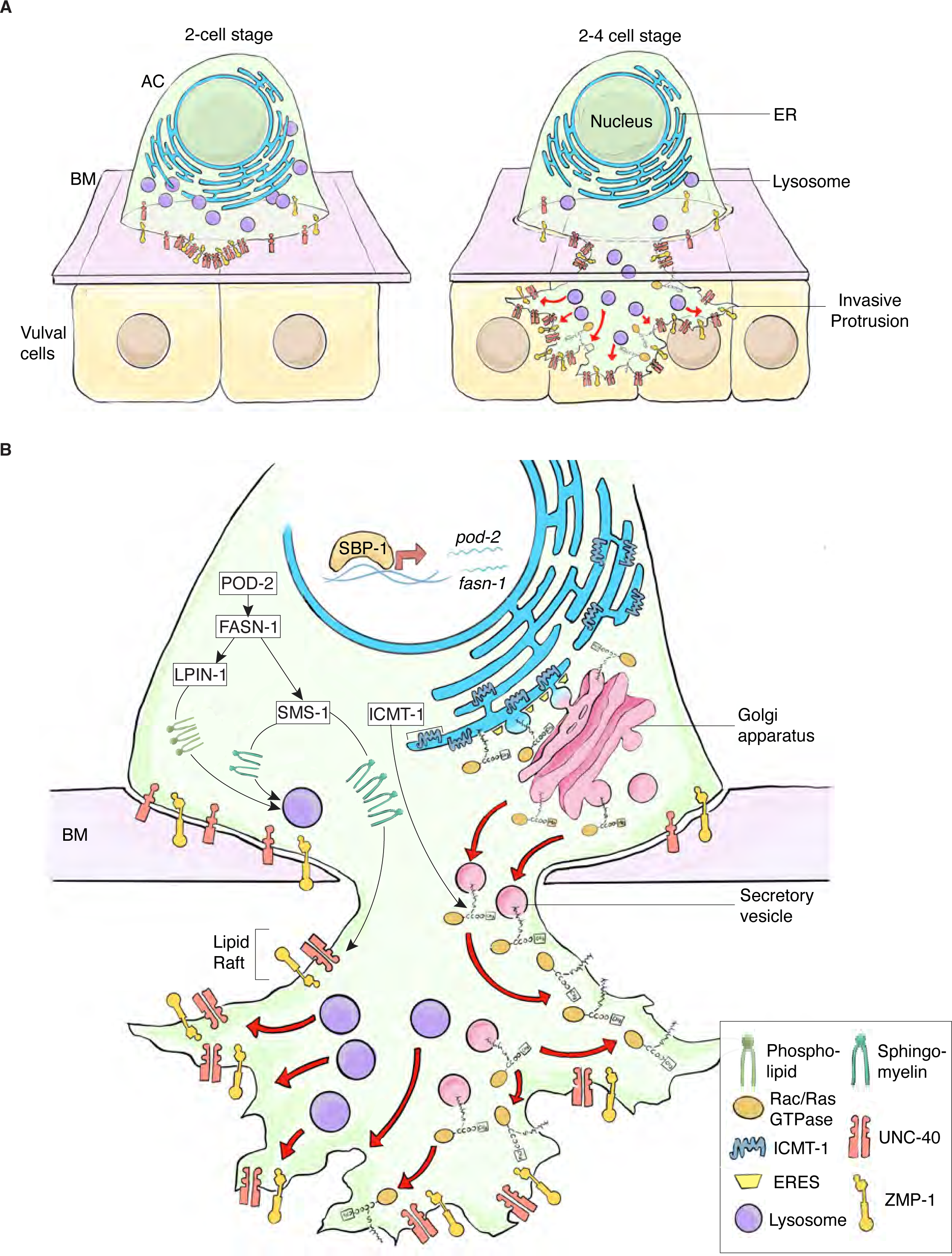
De novo lipid synthesis and polarized prenylation promote invasive protrusion formation. **(A)** A schematic diagram summarizing AC invasion. Invadosomes form and one breaches the underlying BM (P6.p 2-cell stage). The netrin receptor DCC (UNC-40) enriches at the BM breach site and directs lysosomal exocytosis, which provides membrane for protrusion growth (P6.p 2-4-cell stage). AC invasive protrusion growth is further facilitated by the enrichment of GPI-anchored matrix degrading metalloproteinase ZMP-1 and prenylated Rac and Ras-like GTPases that direct F-actin formation. **(B)** A schematic diagram highlighting the roles of de novo lipid synthesis and lipid prenylation during AC invasive protrusion formation. Nuclear SBP-1 promotes expression of the fatty acid synthesis enzymes *pod-2* and *fasn-1* that provide fatty acid substrates for LPIN-1 and SMS-1, which synthesize phospholipids and sphingomyelin that build lysosome stores for the protrusion. Sphingomyelin, which is a key component of lipid rafts, is also required for the enrichment of the UNC-40 receptor and the GPI-anchored metalloprotease ZMP-1, which both partition to lipid rafts. A specialized prenylation system, where the final ER enzyme of prenylation, ICMT-1, localizes to ER exit sites (ERES) that dynamically polarize with the Golgi towards the invasive protrusion, rapidly deliver Rac- and Ras-like GTPases to the AC invasive front to direct protrusion growth.

The invasive protrusion of the AC transiently increases the AC surface area by as much as ∼40% through UNC-40/t-SNARE SNAP-29 mediated lysosome exocytosis (Naegeli et al., 2017). Previous studies have shown that loss of the phosphatidylinositol phosphate kinase *ppk-3*, which regulates the maturation and integrity of lysosomes (Nicot et al., 2006), dramatically reduces the invasive protrusion (Naegeli et al., 2017). While a great deal is known about signaling and transcriptional control of lysosome number and size (Yang and Wang, 2021), the role of lipogenesis in lysosomes is unclear. Our studies indicate that AC lysosome stores are reliant on de novo, cell-autonomous phospholipid synthesis, as AC LPIN-1 levels are highly upregulated and loss of *lpin-1* dramatically reduced AC lysosomal volume prior to invasion, without affecting prenylated Rac GTPase or ZMP-1 (GPI-anchored) localization. Given that phospholipids compose the bulk of membrane lipids (Cockcroft, 2021), this suggests that AC LPIN-1 is likely required to produce phospholipids used to form lysosome stores prior to invasion. Consistent with this notion, we’ve previously reported that the AC ER, where phospholipids are generated, expands dramatically prior to invasion (Cockcroft, 2021; Costa et al., 2023). Loss of the sphingomyelin synthase SMS-1 also reduced AC lysosomes prior to invasion. Although sphingomyelin composes only ∼5% of lipids in most cells, it is made in the Golgi and concentrated in the plasma membrane and within the endolysosome and is important for membrane stability (Goni, 2022; Tang et al., 2022), and thus its loss could affect lysosome stores. Sphingomyelin is also crucial for vesicle biogenesis at the Golgi (Duran et al., 2012), and sphingomyelin reduction possibly reduces the trafficking of phospholipids from the trans-Golgi to the lysosome (Liu et al., 2021). Finally, we discovered that the dolichol branch of the mevalonate pathway is also required for AC lysosome stores. Dolichol phosphate is critical for N-linked protein glycosylation, where dolichols function as a membrane anchor for the transfer of an oligosaccharide to many newly translated proteins within the ER (Esmail and Manolson, 2021; Rauthan and Pilon, 2011). The luminal side of the lysosome membrane contains heavily glycosylated transmembrane proteins, including LAMP-1, LAMP-2, and LIMP-2 (SCAV3), which have been implicated in mediating lysosome integrity by forming a protective glycocalyx that prevents degradation from lysosome hydrolases (Li et al., 2016). Loss of glycosylation after perturbation of the dolichol branch of the mevalonate pathway is thus a strong candidate for the mechanism underlying reduced lysosomes within the AC. Together, these observations reveal the importance of independent lipogenesis pathways in regulating lysosome generation for invasive protrusion formation and suggest targeting lysosome stores might be a promising strategy for limiting cell invasion.

Our results also suggest that the generation of sphingomyelin within the AC is crucial for netrin receptor DCC (UNC-40) localization during the invasive protrusion formation. DCC (UNC-40) localizes to the initial BM breach site, where in response to netrin (UNC-6) secreted by the underlying vulval cells, directs lysosome exocytosis to form the invasive protrusion (Morrissey et al., 2013; Naegeli et al., 2017). Studies in vertebrate commissural neurons and immortalized cells have shown that DCC associates with lipid rafts and lipid raft association is required for netrin mediated activation (Herincs et al., 2005). We found that loss of SMS-1, which generates sphingomyelin, a key component of lipid rafts (Bieberich, 2018; Goni, 2022), perturbed UNC-40 enrichment at the BM breach site. Thus, reduction in the size and growth of the invasive protrusion after loss of SMS-1, could be in part due to perturbation in UNC-40 signaling. Like vertebrates, biochemical studies indicate that *C. elegans* harbor lipid rafts where GPI-anchored proteins partition (Rao et al., 2011; Sangiorgio et al., 2004). Notably, ZMP-1, a GPI-anchored protease, is concentrated in the invasive protrusion. Loss of SMS-1 disrupted ZMP-1 localization, strongly supports the idea that the invasive cell membrane and protrusion are rich in sphingomyelin and lipid rafts that localize and regulate UNC-40, and possibly other proteins that mediate invasion.

Little is known about the regulation of prenylation in cells (Wang and Casey, 2016). Endogenous tagging of the *C. elegans* ER-associated HMG-CoA reductase (HMGR-1) protein of the mevalonate pathway that generates isoprenoids for prenylation, revealed enrichment of HMGR-1 towards the invasive front of the AC. Similarly, the ER-localized proteins of the prenylation branch of the mevalonate pathway, FCE-1 and ICMT-1, which finalize processing of CAAX proteins that have an isoprenoid lipid attached by a prenyltransferase, were also enriched and polarized in the AC. Notably, both HMGR-1 and ICMT-1 were polarized towards the invasive front by UNC-6 (netrin)—an invasive polarity cue generated by the underlying vulval cells that orients components of the invasive machinery, such as the Rac GTPases MIG-2 and CED-10, mitochondria, and the glucose transporter FGT-1 (Wang et al., 2014a; Wang et al., 2014b). These results indicate that polarized prenylation is a part of the cell invasion program. We further discovered that ICMT-1, which is the final enzyme of prenylation (Wang and Casey, 2016), specifically localizes to ER exit sites, which, with the Golgi, dynamically orient towards the invasive protrusion during its formation. Together our results reveal a polarized prenylation and trafficking system, which we found directs the Rac GTPases MIG-2 and CED-10, as well as the Ras-like GTPase RAP-1 robustly to the invasive protrusion. As many GTPases and signaling molecules are prenylated (Borini Etichetti et al., 2020; Rauthan and Pilon, 2011; Wang and Casey, 2016), the AC’s polarized prenylation system might be used to direct many proteins to the invasive protrusion to rapidly direct its formation and functionality.

Only ∼20% of *C. elegans* fatty acids are synthesized de novo, with most being acquired from the diet and imported into cells (Watts and Ristow, 2017). Further, in many organisms, fatty acid synthesis is primarily accomplished by specialized organs, such as the liver, and cells acquire fatty acids from the extracellular environment through lipid import (Cockcroft, 2021). Thus, it is notable that the invasive AC is dependent upon cell-autonomous fatty acid synthesis and uses SREBP (*C. elegans* SBP-1) mediated transcription to direct high levels of expression of the fatty acid synthesis ACC (POD-2) and FASN (FASN-1) enzymes. Further, the AC requires enzymes that mediate phospholipid and sphingolipid synthesis (LPIN-1 and SMS-1) and prenylation promoting enzymes (HMGR-1 and ICMT-1) to generate the invasive protrusion. Numerous studies have implicated the human orthologs of these enzymes in promoting metastasis (Borini Etichetti et al., 2020; Lu et al., 2022; Martin-Perez et al., 2022; Song et al., 2020; Zheng et al., 2019) and pharmacological inhibitors of these enzymes exist (Brohee et al., 2021; Companioni et al., 2021; Marchwicka et al., 2022; Rohrig and Schulze, 2016; Vasseur and Guillaumond, 2022), with HMGCR statin inhibitors showing potential promise in clinical trials targeting invasive glioblastoma in combination with chemotherapy (Afshari et al., 2021). Our results on the requirement of these lipid synthesis and modification enzymes during AC invasion, offers strong in vivo support that using these inhibitors might be an effective strategy to limit the ability of tumors to breach BMs and metastasize.

## Data Availability

*C. elegans* strains and plasmids generated in this study and original data are available upon request. All the other data are included in the manuscript and supplemental materials.

## Acknowledgements

We thank members of the Sherwood laboratory for helpful comments and discussions, P. Schedl for helpful discussions and I. Kenny-Ganzert for making the *lin-29p::mKate2::PLCδ^PH^*construct and the D. Greenstein for DG4324 (*tn1765* (*GFP::3xflag::pod-2*) II). Some strains were provided by the CGC funded by NIH Office of Research Infrastructure Programs (P40 OD010440). We thank the Kiehart laboratory for the use of their NIH funded (R35GM127059) Zeiss Axiovert 200M equipped with a Yokogawa W1 spinning disk confocal head for image collection. This work was supported by the Hargitt fellowship and Jane Coffin Childs Memorial Fund for Medical Research to A.W.J. Soh and R35GM118049 and R21OD032430 to D.R. Sherwood. The authors declare no competing financial interests.

## Author contributions

K. Park, A. Garde, and D.R. Sherwood conceived the project. K. Park designed, conducted all experiments, and analyzed and interpreted the data with the following contributions from other authors: Q. Chi designed and built molecular biology constructs used in Figs. 1, 3-8 and Fig. S3-S5. A. Garde performed the AC invasion RNAi screen and acquired a portion of the AC protrusion time-lapse used in Figs. 2, 4, 5, 6, and Fig. S2. A. Garde also built molecular biology constructs used in Fig. 2 and made the schematics used in Figs. 1, 2, 3, and 6. S.B. Thendral acquired a portion of the AC protrusion time-lapse used in Figs. 2, 4, 5, 6, and Fig. S2. S.B. Thendral also analyzed the AC lysosome volume data used in Figs. 4, 5, 6, and Fig. S4 and made the schematics used in Figs. 5, 8, and 9. A.W.J. Soh analyzed the AC protrusion time-lapse data used in Figs. 2, 4, 5, 6, and Fig. S2. A.W.J. Soh also analyzed the GTPase and matrix metalloproteinase polarization data used in Figs. 4, 5, and 6. S.B. K. Park prepared all figures. D. R. Sherwood, K. Park, and A. Garde wrote the manuscript. D. R. Sherwood, K. Park, A. Garde, A.W.J. Soh, and S.B. Thendral edited the manuscript.

## Materials and methods

### Strains and culture conditions

*C. elegans* strains were maintained on nematode growth medium (NGM) plates and fed with *E. coli* (OP50) at 16-20°C following standard worm growth conditions (Stiernagle, 2006). Please refer to Table S3 for strains used in this study. In this manuscript, we designate a linkage between the promoter (p) and the open reading frame with a double semicolon (::). Strains sourced from Caenorhabditis Genetics Center (CGC) are to be requested directly from CGC. Further requests for reagents should be directed to the corresponding author, D.R. Sherwood.

### Construction of genome edited strains

Fluorescently tagged strains were generated using CRISPR-Cas9-mediated genome editing as previously described (Dickinson et al., 2013; Keeley et al., 2020). Briefly, *mNeonGreen (mNG)* or *mScarlet* coding sequence was knocked into a gene of interest via CRISPR-Cas9-triggered homologous recombination using a short guide RNA (sgRNA) and a homologous repair template that contained the fluorophore coding sequence and selection markers. Silent mutations were introduced into the homology arms using site-directed mutagenesis (refer to Table S4 for information on sgRNA sequences). All sgRNAs were cloned into the pDD122 Cas9-sgRNA expression vector (Table S5; (Dickinson et al., 2013)). The Cas9-sgRNA plasmid and homologous repair template were co-injected into the gonad of young adult N2 hermaphrodites. Recombinant animals were identified in the F3 offspring of injected animals based on the presence of selectable markers (hygromycin (Table S5) resistance and dominant-negative *sqt-1 rol* phenotype). Selection markers were subsequently excised from the genome through Cre-Lox recombination. Proper genome editing was confirmed by DNA sequencing of the edited genomic locus and health of the genome edited line determined as previously described (Keeley et al., 2020). Please refer to Table S4 for information on genotyping primers.

### Construction of fusion proteins

To express ICMT-1 in the AC, we used the *cdh-3* promoter (Garde et al., 2022) and generated *cdh-3p::icmt-1::mScarlet* by PCR fusion. Specifically, a 1.1 kb fragment of the full length *icmt-1* sequence was fused with a 1.5 kb fragment of the *cdh-3* promoter and *mScarlet* coding region in the pAP088 vector (Table S5; (Pani and Goldstein, 2018)). The same strategy was applied using a 1.5 kb fragment of *lin-29* promoter in the pAP088 vector (*lin-29p::icmt-1::mScarlet)*. To construct the AC-specific lysosome marker *cdh-3p::lmp-1::mNG* using PCR fusion, a 0.9 kb fragment of the full length *lmp-1* sequence was fused with a 1.5 kb fragment of the *cdh3* promoter and *mNG* region in the pAP088 vector. To build the AC-specific membrane marker *lin-29p::mKate2::PH*, a 1.5 kb fragment of *lin-29* promoter was fused with *mKate2* and 0.5kbp of the *PLCδ^PH^* in the pAP088 vector. Refer to Table S4 for primer sequences. All constructs were co-injected into the gonad of young adult N2 hermaphrodites with the Cas9-sgRNA plasmid (pCFJ352; Table S5). Single-copy transgenes were knocked-in at the ttTi4348 (I) locus in chromosome I using the pAP088 vector (Frokjaer-Jensen et al., 2012). Recombinant animals were identified in the F3 offspring of P0 injected animals based on the presence of selectable markers (hygromycin resistance and dominant-negative *sqt-1 rol* phenotype). Selection markers were subsequently excised from the genome through Cre-Lox recombination.

### Construction of RNAi plasmids

RNAi constructs targeting *sbp-1*, *hmgr-1*, *icmt-1*, and *B0024.13* were obtained from the Ahringer laboratory and Vidal laboratory RNAi libraries (Kamath and Ahringer, 2003; Rual et al., 2004). To improve the efficiency of RNAi knockdown targeting *sbp-1*, *hmgr-1*, *icmt-1*, and *B0024.13*, new RNAi constructs were synthesized. These constructs targeted the entire mRNA transcripts of *sbp-1*, *hmgr-1*, *icmt-1*, and *B0024.13* and were designed using the T444T RNAi vector (Sturm et al., 2018). The T444T vector was digested using SacII and HindIII restriction enzymes and ligated with the PCR-amplified coding sequences via Gibson assembly. Please refer to Table S4 for primer sequences.

### RNAi Experiments

Bacteria harboring the RNAi vectors were cultured in selective media (lysogeny broth with 100 mg/ml ampicillin) for 16 h at 37°C. Transcription of double-stranded RNAi was induced with 1mM Isopropyl ß-D-1-thiogalactopyranoside (IPTG) for one h at 37°C. Bacteria cultures were plated onto NGM plates that were treated with 1 mM IPTG and 100 mg/ml ampicillin. Plates were dried at room temperature for 12 h before use. RNAi experiments were conducted by feeding L1 worms with HT115(DE3) bacteria expressing double-stranded RNAi for 36-42 h at 18°C or 20°C. Bacteria expressing the empty RNAi vector (L4440 or T444T) served as the negative control in all experiments. Whole animal RNAi knockdown was performed in this study unless otherwise specified. AC-specific RNAi experiments were performed using the strain NK1316 (Garde et al., 2022). This strain harbors a loss of function mutation in *rde-1*(*ne219*), an argonaute protein required for RNAi. Expression of *rde-1* in the AC (*fos-1ap::rde-1*) restores RNAi specifically in the AC. The RNAi screen was performed to identify lipid synthesis enzymes (Table S2) that are required for AC invasion (Table S2). For enzymes with multiple family members (*i.e.,* SMS-1, −2, −3, CGT-1, −2, −3, and PSSY-1, −2), the enzyme with expression in the AC as determined by previously published AC-specific RNA-Seq (Costa et al., 2023) was examined in the RNAi screen. To enhance RNAi efficiency, two strategies were used. RNAi was performed in the *rrf-3*(*pk1426*) background, which harbors a mutation in an RNA-directed RNA polymerase whose loss sensitizes the worms to RNAi (Costa et al., 2023; Simmer et al., 2002), or performed transgenerationally. RNAi knockdown efficiencies were determined using endogenously tagged fluorescent protein strains as described in the ‘Quantification and statistical analysis’ section and reported in Table S1.

### Microscopy and image acquisition

The AC invasion RNAi screen was performed using an Axio Imager A1 microscope (Carl Zeiss), which was equipped with a Plan-APOCHROMAT 100x (1.4 NA) differential interference contrast (DIC) oil immersion objective. Images were acquired with a Zeiss AxioCam 305 mono CMOS camera controlled by the Zen Blue version 3.2 software (Carl Zeiss). For enhanced image resolution of ER-resident proteins FCE-1, HMGR-1, and ICMT-1, imaging was performed on the Zeiss Axiovert 200M microscope using a Zeiss Plan-Apochromat 63X (1.4 NA) oil immersion objective, which was mounted with a CSU-W1 spinning disc confocal (Yokogawa). Images were acquired with an Orca-Fusion BT camera (Hamamatsu Photonics) controlled by µmanager software (Edelstein et al., 2010). For single time point AC snapshots to visualize fluorescently tagged proteins in the AC, worms were anesthetized in 0.01M sodium azide and mounted on 5% noble agar pad with a #1.5 cover slip. Fluorescence images were acquired as confocal z-stacks spanning the entire cell (total z-range: 5-11 µm) using 0.37 µm optical z-slice. For low abundance proteins that are prone to photobleaching, a shorter total z-range (5-7 µm) and larger optical z-slice (0.50 µm) were used. Imaging experiments were performed using an Axio Imager A1 microscope (Carl Zeiss) equipped with a CSU-10 spinning disc confocal (Yokogawa) and a Plan-APOCHROMAT 100x (1.4 NA) oil immersion objective. Images were acquired using either a Hamamatsu Orca-Fusion sCMOS camera or ImagEM EM-CCD camera that was controlled by µmanager software (Edelstein et al., 2010). For time-lapse imaging, worms were anesthetized in 5 mM of levamisole and mounted on 5% noble agar pad with a #1.5 cover slip. VALAP was used to enclose samples to prevent evaporation. Each live imaging experiment was limited to under 2 h to prevent starvation induced changes in physiology (Schindler et al., 2014). To track the growth of AC invasive protrusion, confocal z-stacks (total z-range: 8-11µm; optical z-slice: 0.37 µm) of control animals were acquired from the late 2-cell stage, when the protrusion forms, until the mid 4-cell stage when the protrusion retracts (total imaging duration ∼90 min). Due to the AC delay in BM breaching after RNAi mediated reduction of s*bp-1*, *lpin-1*, *sms-1*, and *hmgr-1*, RNAi-treated animals were imaged beginning at the P6.p 2-4 cell stage to capture protrusion formation. The imaging frame interval for all movies was 5 min except for *sms-1* RNAi-treatment and its corresponding control, which were imaged at 4 min intervals. To visualize UNC-40 recruitment to the invadosome site of BM breach, worms were manually rolled to align the ventral surface along the imaging plane as previously described (Kelley et al., 2017). An imaging frame interval of 60 sec was used. All image acquisition was carried out at 20°C.

### Quantification and statistical analysis

#### RNAi knockdown efficiency

The efficiency of RNAi knockdown was assessed by quantifying the fluorescence intensity of each fluorophore-tagged protein of interest (mNG::SBP-1, mNG::HMGR-1, mNG::LPIN-1, SMS-1::mNG, FNTA-1::mNG, and ICMT-1::mScarlet) in empty RNAi vector control and RNAi gene targeted knockdown animals. All z-stack images were sum intensity z-projected (total z-range: 0.74–1.48µm) except for FNTA-1::mNG and ICMT-1::mScarlet whereby only a single z-slice that spans across the AC longitudinally was acquired. The mean fluorescence intensity of each targeted protein in the AC was quantified by drawing an AC outline with the “Freehand selection” tool and computed via Fiji 1.53f (Schroeder et al., 2021). To account for and subtract autofluorescence signals, all fluorescence intensity comparisons were made against wild-type animals (N2). The local background mean fluorescence intensity was quantified from the adjacent uterine tissue region where autofluorescence signals were observed. The net mean fluorescence intensity was then calculated by subtracting the local background mean fluorescence intensity from the AC mean fluorescence intensity in control and RNAi knockdown animals.

#### Scoring of AC Invasion

AC invasion was assessed via DIC microscopy as previously described (Sherwood et al., 2005). Basement membrane (BM) breach was confirmed by fluorescence microscopy using a fluorescently tagged BM laminin marker (LAM-1::GFP; NK1316). Scoring was performed at the 1° VPC P6.p 4-cell stage when the clearance of the BM is completed in wild-type animals. Invasion was scored as “full invasion” if the BM breach was at least half of the AC width. Invasion was scored as “partial invasion” if the BM breach was at most one-third of the AC width. Invasion was scored as “blocked invasion” if the BM underneath the AC was completely intact. Both partial and blocked invasion were categorized as an invasion defect.

#### Quantification of AC invasive protrusion volume

AC invasive protrusion volume was quantified as previously described (Kelley et al., 2017; Naegeli et al., 2017). Confocal microscopy was performed to capture z-stacks (total z-range: 8-11µm; 0.37µm optical slice) of the AC (mCherry::PLCδ^PH^) and the underlying BM (LAM-1::GFP). Confocal z-stacks were used to make 3D reconstructions of the AC and the BM using Imaris 9.9 (Bitplane; (Naegeli et al., 2017)). Isosurface renderings of mCherry::PLCδ^PH^ were created by empirically determined fluorescence intensity thresholds that outline the AC plasma membrane. To distinguish the AC invasive protrusion from the rest of the cell, the Imaris isosurface slicer tool was used to cut the AC isosurface at the AC-BM junction. AC invasive protrusion was defined by the region of the AC that breached the BM. AC protrusion volume was tracked over a 60 min period (images acquired every 5 min) in control animals and *sbp-1*, *lpin-1*, and *hmgr-1* RNAi treated animals beginning at the time when protrusion formation first occurred. In all cases the protrusion retracted or failed to grow further over by the final 15 min of analysis, which ensured that maximum protrusion volume was observed. Notably, in two control animals, protrusion formation initiated earlier, and these were included in the analysis (see Fig. S2 B). For *sms-1* RNAi treated animals and their corresponding controls, the AC protrusion volume was tracked for 64 min (images acquired ever 4 min). The AC invasive protrusion volume was quantified using the Imaris volume measurement tool in control and RNAi treated animals.

#### Quantification of AC lysosome volume

3-dimensional reconstructions were created from confocal z-stacks of the AC expressing *lin-29p*::LMP-1::mNG using Imaris 9.9 (Bitplane; (Naegeli et al., 2017)). Due to the variability in lysosome fluorescence signals, a fluorescence intensity threshold was determined empirically for each AC to distinguish lysosomes from the background fluorescence signals. Isosurface renderings of LMP-1::mNG were then created and the total AC lysosomal volume was quantified using the Imaris volume measurement tool in control and RNAi treated animals.

#### Quantification of UNC-40 levels at the AC BM breach site

To visualize UNC-40 at the BM breach site, animals expressing UNC-40::GFP and the BM marker LAM-1::mCherry were manually rolled to align the ventral surface along the imaging plane and then imaged by confocal microscopy. Confocal z-stacks that span the BM breach and the AC invasive membrane (total z-range: 5µm; optical z-slice: 0.37µm) were acquired. To visualize UNC-40::GFP localization at the BM breach, a subset of z-stack images (z-range: 1.48µm) spanning across the BM breach were sum projected using the “Z Projection Sum Slices tool” in Fiji 1.53f (Schroeder et al., 2021). The mean fluorescence intensity of UNC-40::GFP at the BM breach site was quantified using the Fiji “Measure” tool and the mean fluorescence intensity outside the worm was measured to account for background. The net mean fluorescence intensity was then calculated by subtracting the background mean fluorescence intensity from the AC mean fluorescence intensity in control and RNAi knockdown animals. UNC-40 localization was assessed at the P6.p 2-cell stage in control animals. Due to the AC invasion delay in *sms-1* RNAi treated animals, UNC-40 localization was assessed from the P6.p 2-4-cell stage to the 4-cell stage.

#### Quantification of GTPases and ZMP-1 polarization at the AC invasive membrane

Enrichment of GTPases (GFP::CED-10, GFP::MIG-2, mNG::RAP-1), GFP::CAAX, and the matrix metalloproteinase ZMP-1 (ZMP-1::mNG) was assessed using Fiji 1.53f (Schroeder et al., 2021). To reduce out-of-focus fluorescence signals from neighboring tissues that were positioned above and below the AC z-plane, image sub-stacks that only span across the AC center were sum intensity z-projected (total z-range: 1.48µm; optical z-slice: 0.37µm). Next, the mean fluorescence intensity of each protein at the basal and apical AC membrane was quantified with a 5-pixel-wide line using the Fiji ‘Freehand selection’ tool (Wang et al., 2014a). The local background mean fluorescence intensity outside the worm was measured and subtracted from the apical and basal membrane mean fluorescence intensities. GTPase, CAAX::GFP, and ZMP-1 polarization was then determined by calculating the ratio of the basal/apical fluorescence intensities.

#### Assessment of the ER prenylation enzyme ICMT-1 colocalization with HMGR-1 and FCE-1

To assess ER proteins’ colocalization, confocal z-stacks of fluorescently co-labelled strains (ICMT-1::mScarlet and mNG::HMGR-1, ICMT-1::mScarlet and mNG::FCE-1) were acquired. To visualize the AC ER, the centermost AC z-slice was selected. To visualize the ER proteins clearly, Gaussian Blur (1 pixel) was applied Background mean fluorescence intensity was accounted for by subtracting the mean fluorescence intensity outside the worm from the respective fluorescence channel. A 5-pixel-wide line was then drawn to quantify the relative fluorescence intensity distribution along this line. The intensity profile for each fluorescently tagged protein was normalized to its maximum fluorescence intensity.

#### Analysis of the AC ER exit site and Golgi polarization

To quantify ER exit site (mNG::SEC-16A.1) polarization in the AC, 3-dimensional reconstructions were created from confocal z-stacks of the AC expressing mNG::SEC-16A.1 and the AC specific plasma membrane marker *lin-29p*::mKate2::PLCδ^PH^ using Imaris 9.9 (Bitplane). Isosurface renderings of mKate2::PLCδ^PH^ were created by determining fluorescence intensity thresholds that outline the AC plasma membrane. mNG::SEC-16A.1 punctae number in the AC basal and apical halves were counted and expressed as basal/apical ratio to assess mNG::SEC-16A.1 polarization. To quantify Golgi polarization in the AC, 3-dimensional reconstructions were created from confocal z-stacks of the AC expressing the Golgi marker AMAN-2::GFP (*cdh-3p*::AMAN-2::GFP) using Imaris 9.9 (Bitplane). AMAN-2::GFP punctae number in the AC basal and apical halves were counted and expressed as basal/apical ratio to assess Golgi polarization.

#### Assessment of the AC ER exit site colocalization with the prenylation enzyme ICMT-1

To assess colocalization between the ER exit site (mNG::SEC-16A.1) and the ER-resident prenylation enzyme (ICMT-1), 3-dimensional reconstructions were created from confocal z-stacks of endogenously tagged mNG::SEC-16A.1 and AC-expressed ICMT-1::mScarlet (*lin-29p::icmt-1::mScarlet*) using Imaris 9.9 software (Bitplane). The percentage of mNG::SEC-16A.1 punctae that overlapped with AC-expressed ICMT-1::mScarlet punctae in the AC was then determined.

#### Statistical analysis

All experiments were performed with two or more replicates for both control and experimental results. Statistical analysis was performed using GraphPad Prism 10. Normality of data distribution was evaluated with the D’Agostino-Pearson normality test. Student’s t-test was performed on samples with Gaussian distribution. Mann-Whitney U test was performed on samples with non-Gaussian distribution. For comparisons of three or more datasets, One-way ANOVA followed by Brown Forsythe and Welch ANOVA tests (samples with Gaussian distribution) or Kruskal-Wallis test and Dunn’s multiple comparisons test (samples with non-Gaussian distribution) was performed. To compare categorical data Fisher’s exact test was performed.

## Online Supplemental Material

Fig. S1, related to Figs. 2-9, shows the substrates and enzymes involved in de novo fatty acid, phospholipid and sphingolipid synthesis and mevalonate pathways in *C. elegans*. Fig. S2, related to Figs. 2, 4, 5, and 6, describes the AC invasive protrusion formation on control and corresponding RNAi-targeted lipid synthesis and prenylation genes (*sbp-1*, *lpin-1*, *sms-1*, and *hmgr-1*). Fig. S3, related to Fig. 3, shows the expression of phospholipid and sphingomyelin synthesis genes (*pod-2, fasn-1*, *elo-1*, *lpin-1*, and *sms-1*) required for AC invasion. Fig. S4, related to Figs. 6 and 7, shows the regulation of AC lysosomal volume and GTPase polarization regulation by the mevalonate pathway enzymes (ICMT-1 and B0024.13), and independence of *hmgr-1* expression from SBP-1. Fig. S5, related to Figs 6-8, shows UNC-6 (netrin) directed polarization of the AC prenylation enzymes (HMGR-1 and ICMT-1) and time course of AC GTPase polarization at the invasive front. Table S1, related to Figs. 3-6, and Figs. S4 and S5, shows RNAi knockdown efficiency for respective experiments. Table S2, related to Figs. 2-9 and Figs. S1-S5, shows RNAi knockdown and gene mutations effects on AC invasion. Table S3 shows the primer sequences used to generate genome-edited strains, transgenic constructs, and RNAi plasmids. Table S4 outlines the strains used in this study and Table S5 lists the chemical and DNA reagents used.

---

**Figure S1.**
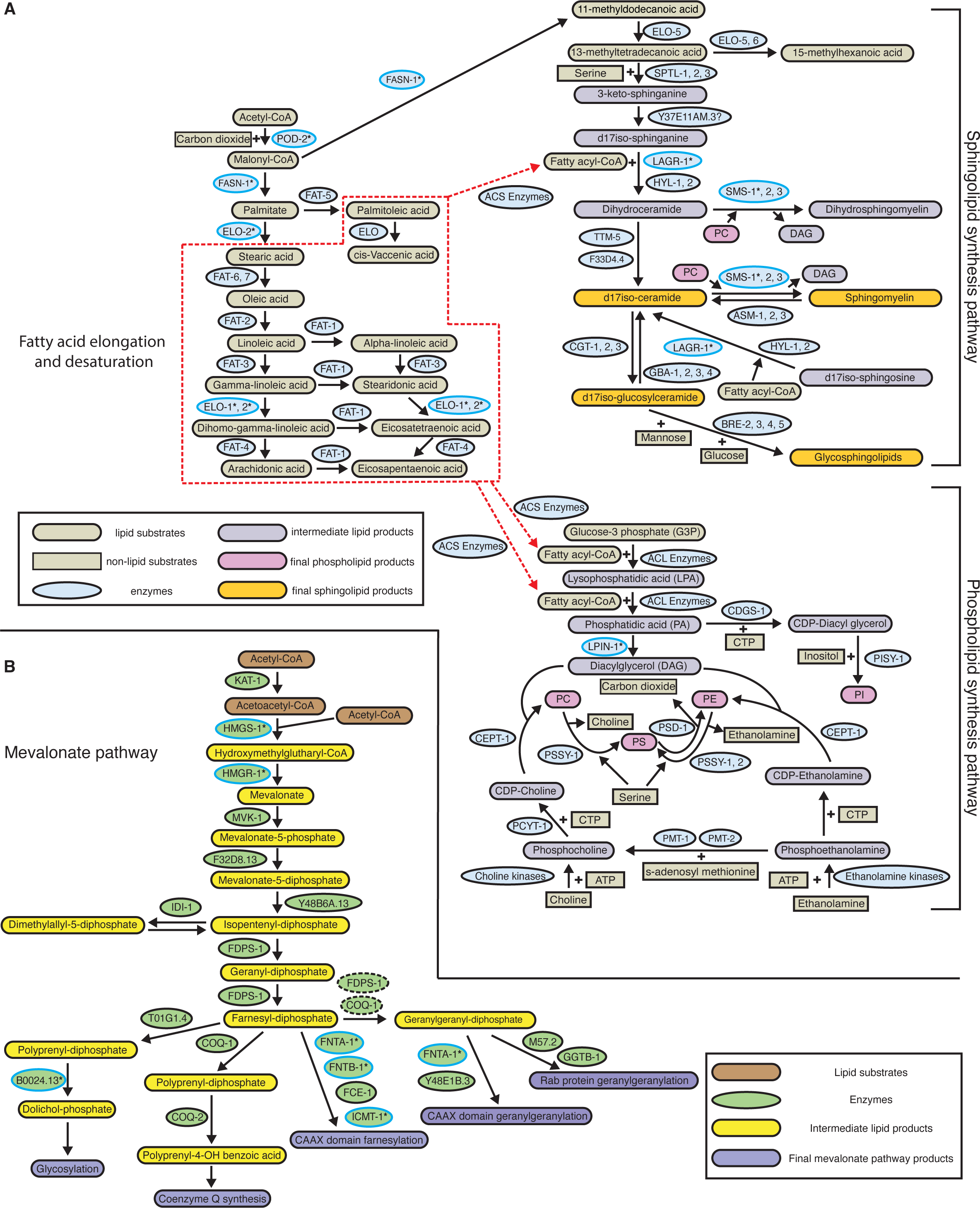
Fatty acid, phospholipid, and sphingolipid synthesis and the mevalonate pathways in *C. elegans*. **(A)** A schematic diagram shows the enzymes and substrates that are involved in processing acetyl-CoA (derived from glucose/mitochondrial citrate) to provide fatty acyl-CoA for the phospholipid and sphingolipid synthesis pathways. For reactions catalyzed by multiple enzymes with redundant functions (SMS-1, 2, 3, CGT-1, 2, 3, and PSSY-1, 2), the enzyme with expression in the AC as determined by AC-specific RNA-Seq was examined in the RNAi screen (Table S2, see Methods). The enzymes whose RNAi loss led to a significant invasion defect are marked by asterisks (refer to Table S2). ACS, Acyl-CoA synthetase, represents 23 *C. elegans* homologs with predicted roles in Acyl-CoA synthesis; ACL, Acyl-CoA ligase, represents 14 *C. elegans* homologs with predicted activity as acyl-transferases; ATP, adenosine triphosphate; CTP, cytidine triphosphate; PC, phosphatidylcholine; PE, phosphatidylethanolamine; PI, phosphatidylinositol; PS, phosphatidylserine. **(B)** A schematic diagram showing the enzymes and substrates of the mevalonate pathway that promote Rab and CAAX geranylgeranylation, CAAX farnesylation, coenzyme Q synthesis, and protein glycosylation. The enzymes whose RNAi loss led to a significant invasion defect are marked by asterisks (refer to Table S2). COQ-1 and FDPS-1 in green dashed circle are the closest known homologs to the human geranylgeranyl diphosphate synthase. Schematics were adapted from Watts and Ristow 2017, Genetics. 207(2):413-446.

**Figure S2.**
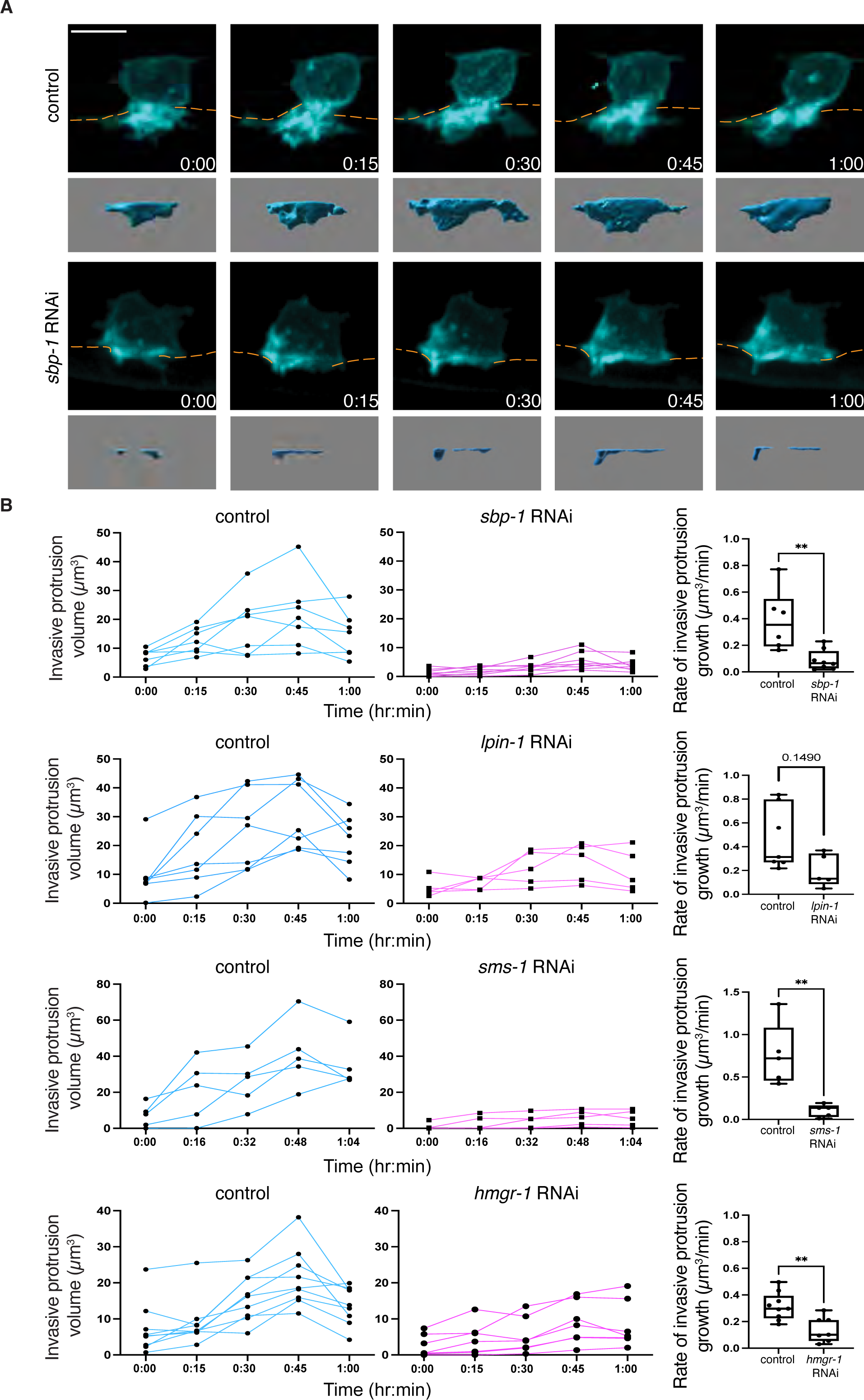
SBP-1, LPIN-1, SMS-1, and HMGR-1 promote AC invasive protrusion growth. **(A)** Maximum intensity z-projected timelapse fluorescence images showing the formation of the AC invasive protrusion (mCherry::PLCδ^PH^, cyan) in a control and *sbp-1* RNAi knockdown animal (BM, orange dashed lines). Time points indicate h::min. Frame interval, 15 min. Isosurfaces of the AC invasive protrusions are shown below. The AC invasive protrusion after *lpin-1*, *sms-1*, and *hmgr-1* RNAi is shown in Figs. 4-6 and Videos 2-4. **(B)** (*Left*) Line graphs of invasive protrusion volume over time in control (cyan) animals and in animals after *sbp-1*, *lpin-1*, *sms-1*, and *hmgr-1* AC-specific RNAi treatment (magenta). (Right) Boxplots of the AC invasive protrusion growth rates. In this and subsequent figures, box edges indicate the 25^th^ and 75^th^ percentiles, whiskers the maximum and minimum values, and the line inside each box the median value. AC protrusion growth rate over 60 or 64 min was calculated (n ≥ 5 animals per condition, ** p ≤ 0.01, p = 0.1490, Mann-Whitney U test). All data are from two or more replicates. Scale bar, 5µm.

**Figure S3.**
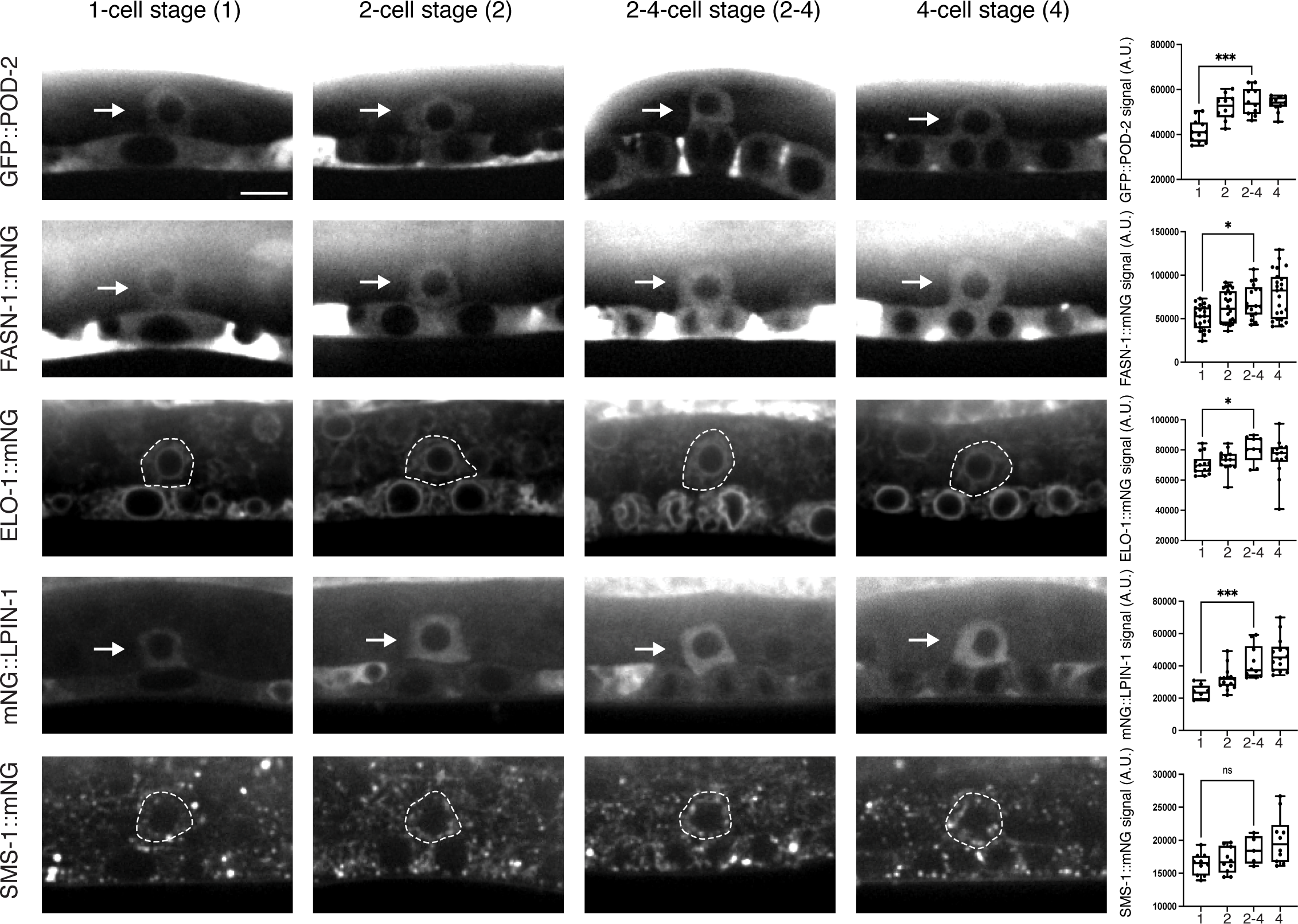
POD-2, FASN-1, ELO-1, LPIN-1, and SMS-1 AC expression from P6.p 1-to 4-cell stage. (*Left*) Sum intensity z-projected fluorescence images of endogenously tagged GFP::POD-2, FASN-1::mNG, ELO-1::mNG, mNG::LPIN-1 and SMS-1::mNG in the AC (arrows and white dotted lines) from the P6.p 1-cell to 4-cell stage. (*Right)* Boxplots show the mean fluorescence intensity of each protein in the AC (n ≥ 5 animals per condition * p ≤ 0.05, *** p ≤ 0.001, ns (not statistically significant), p > 0.05, One-way ANOVA followed by Brown Forsythe and Welch ANOVA tests and One-way ANOVA followed by Kruskal-Wallis test and Dunn’s multiple comparisons test). All data are from two or more replicates. Scale bar, 5µm.

**Figure S4.**
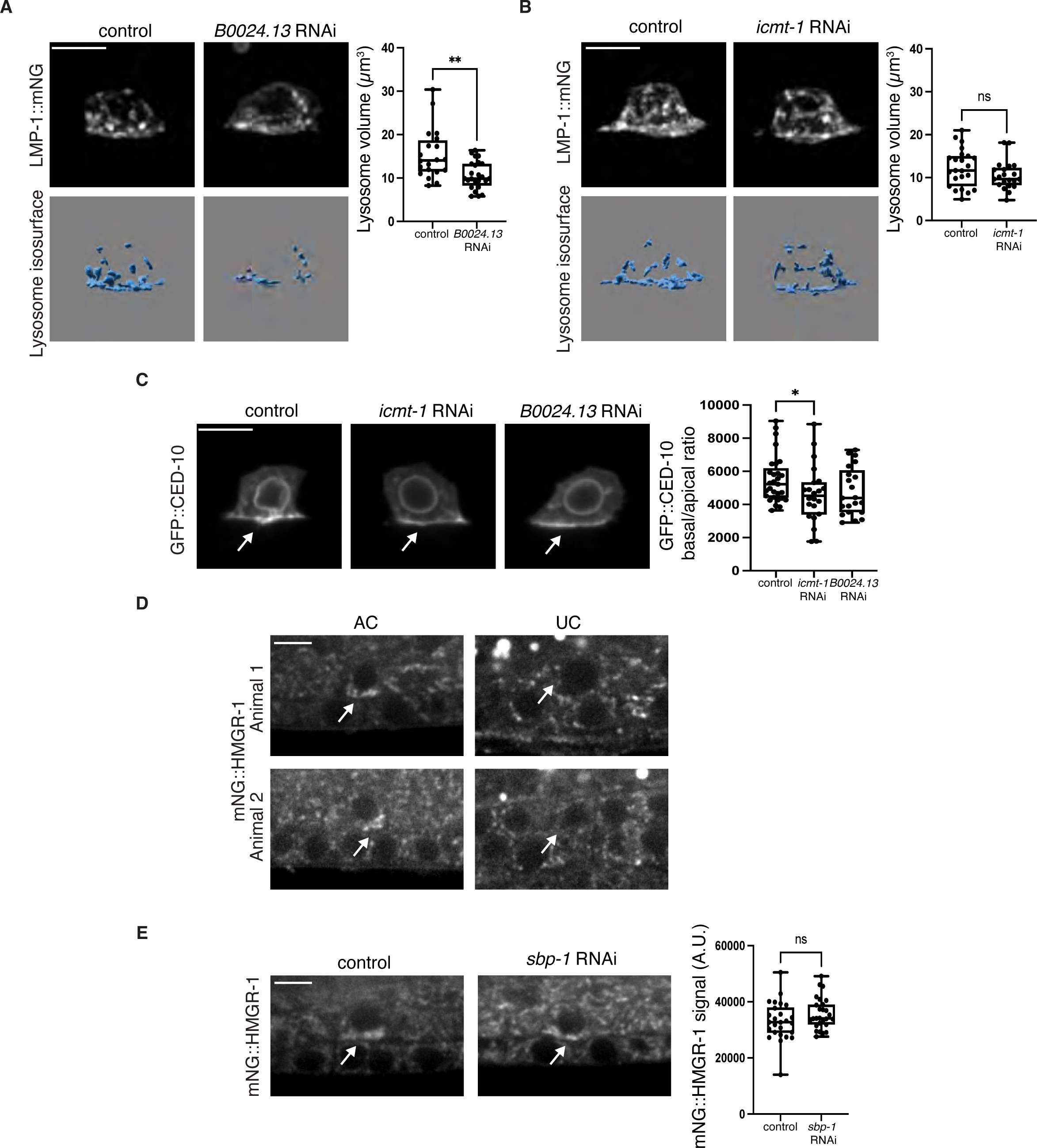
Role and regulation of mevalonate pathway in invasive protrusion formation. **(A)** (*Left*) Sum intensity z-projected fluorescence and isosurface images showing the AC lysosomes (LMP-1::mNG) in a control and in a *B0024.13* (dolichol synthesis) RNAi treated animal at the initiation of invasive protrusion formation. (*Right*) Boxplot of AC lysosomal volume in control and *B0024.13* RNAi treated animals (n ≥ 20 animals per condition, ** p ≤ 0.01, Mann-Whitney U test). **(B)** Sum intensity z-projected fluorescence and isosurface images showing the AC lysosomes (LMP-1::mNG) in a control and *icmt-1* RNAi treated animal at the initiation of invasive protrusion formation. Boxplot of AC lysosome volume in control and *icmt-1* RNAi treated animals (n ≥ 21 animals per condition. ns (not statistically significant), p > 0.05, unpaired two-tailed Student’s t-test). **(C)** (*Left*) Sum intensity z-projection fluorescence images of GFP::CED-10 shows enrichment at the AC basal plasma membrane (arrow) in a control, an *icmt-1*, and a *B0024.13* RNAi treated animal at the initiation of invasive protrusion formation. (*Right)* Boxplot showing the basal/apical ratio of GFP::CED-10 fluorescence intensity (n ≥ 21 animals per condition, * p ≤ 0.05, Mann-Whitney U test). **(D)** (*Left*) Sum intensity z-projected fluorescence images of mNG::HMGR-1 in two ACs with mNG::HMGR-1 enriched at the invasive basal side (arrows). (*Right*) mNG::HMGR-1 is not concentrated at the basal side of uterine cells (UC). **(E)** (*Left*) Sum intensity z-projected fluorescence images of mNG::HMGR-1 in the AC (arrows show AC and enrichment at invasive front) in a control and a *sbp-1* RNAi treated animal. (*Right*) Boxplot of mNG::HMGR-1 fluorescence intensity in the AC of control and *sbp-1* RNAi animals (n ≥ 26 animals per condition, unpaired two-tailed Student’s t-test, ns (not statistically significant), p > 0.05, unpaired two-tailed Student’s t-test). All data are from two or more replicates. Scale bars, 5µm.

**Figure S5.**
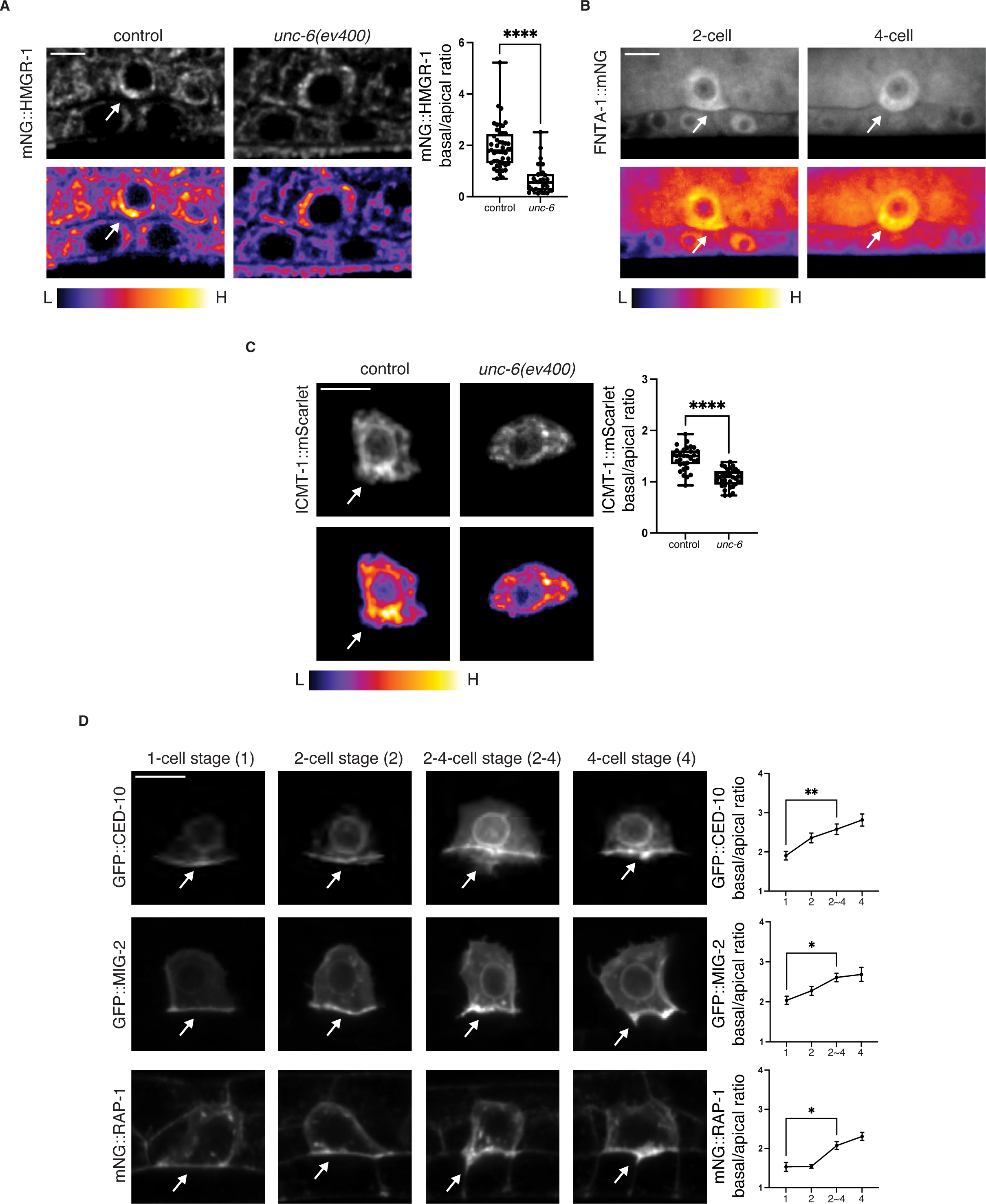
Regulation and expression of prenylation enzymes and GTPases during AC.invasion. **(A)** (*Left*) Sum intensity z-projected fluorescence images of mNG::HMGR-1 in a wild-type and an *unc-6*(*ev400*) mutant animal at the initiation of invasive protrusion formation. Bottom panels show spectral fluorescence-intensity maps, which display the minimum (L, low) and maximum (H, high) pixel value range of the acquired data. (*Right*) Boxplot shows AC basal/apical ratio of mNG::HMGR-1 fluorescence intensity in wild-type and *unc-6*(*ev400*) animals (n ≥ 30 animals per condition, **** p ≤ 0.0001, Mann-Whitney U test). **(B)** Sum intensity z-projected images show high levels of FNTA-1::mNG in the AC at the P6.p 2-cell stage (n = 12/12 animals) and 4-cell (n = 12/12 animals) stage. FNTA-1::mNG basal enrichment was often observed at the P6.p 4-cell stage (n = 12/12 animals). Bottom panels show spectral fluorescence-intensity maps, which display the minimum (L, low) and maximum (H, high) pixel value range of the acquired data. **(C)** Sum intensity z-projected images of ICMT-1::mScarlet in the AC of a wild-type and an *unc-6*(*ev400*) mutant animal at the initiation of invasive protrusion formation. Bottom panels show spectral fluorescence-intensity maps, which display the minimum (L, low) and maximum (H, high) pixel value range of the acquired data. (*Right*) Boxplot shows AC basal/apical ratio of ICMT-1::mScarlet in wild type and *unc-6*(*ev400*) animals (n ≥ 31 animals per condition, **** p ≤ 0.0001, unpaired two-tailed Student’s t-test). **(D)** (*Left*) Sum intensity z-projected images showing the time course of GFP::CED-10, GFP::MIG-2, and mNG::RAP-1 polarization from the P6.p 1-cell to 4-cell stage. *(Right)* Line graphs of AC basal/apical polarization of each protein’s fluorescence intensity (n ≥ 10 animals per condition, * p ≤ 0.05, ** p ≤ 0.01, One-way ANOVA followed by Brown Forsythe and Welch ANOVA tests and One-way ANOVA followed by Kruskal-Wallis test and Dunn’s multiple comparisons test). All data are from two or more replicates. Scale bars, 5µm.

**Table S1.** RNAi knockdown efficiency.

| Gene targeted | Strain genotype<sup>a</sup> | Treatment<sup>b</sup> | Percent protein knockdown (%) | n<sup>c</sup> |
| --- | --- | --- | --- | --- |
| *sbp-1* | *qy94[mNG::sbp-1]* III | *sbp-1* (RNAi) (T) | 73.7 | ≥11 |
| *lpin-1* | *qy208[mNG::lpin-1]* V | *lpin-1* (RNAi) (L) | 95.1 | ≥12 |
| *hmgr-1* | *qy202[mNG::hmgr-1]* III | *hmgr-1* (RNAi) (L) | 99.3 | 15 |
| *hmgr-1* | *qy202[mNG::hmgr-1]* III | *hmgr-1* (RNAi) (T) | 100 | ≥19 |
| *sms-1* | *rrf-3(pk1426)* II; *qy198[sms-1::mNG]* IV | *sms-1* (RNAi) (L) | 61.9 | ≥22 |
| *fnta-1* | *qy199[fnta-1::mNG]* IV | *fnta-1* (RNAi) (L) | 85 | ≥11 |
| *icmt-1* | *qy205[lin-29p::icmt-1::mscarlet]* I | *icmt-1* (RNAi) (T) | 71.8 | ≥13 |

<sup>a</sup> All RNAi was performed using strains endogenously edited with fluorophores with the exception of *sms-1* RNAi, which was conducted using a strain harboring *rrf-3* (*pk1426*) mutation that sensitizes the strain to RNAi.
<sup>b</sup> L stands for L4440 vector, and T stands for T444T vector.
<sup>c</sup> Number of animals imaged in each condition

**Table S2.**
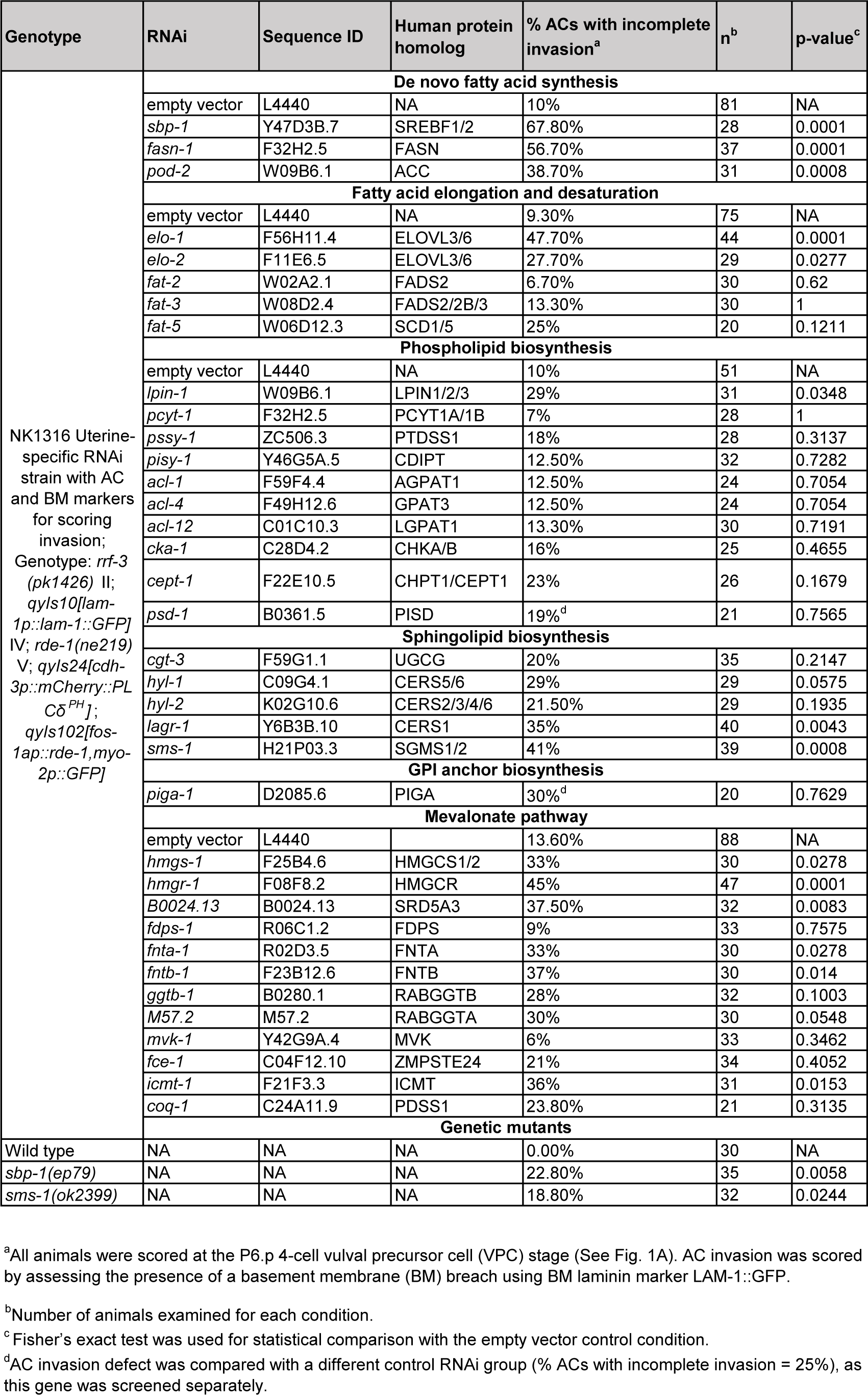
Lipogenesis and lipid modification AC invasion screen.

**Table S3.**
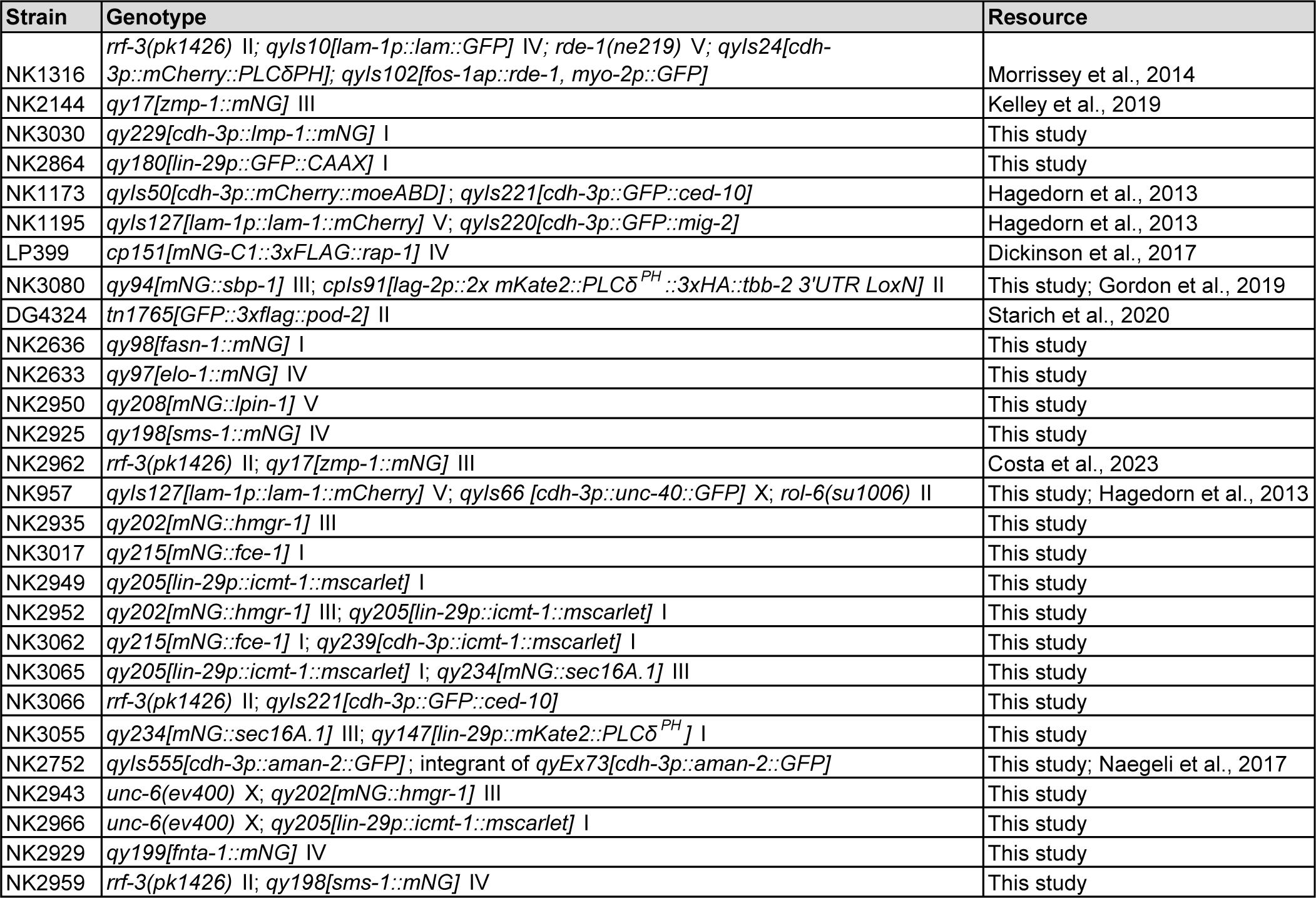
*C. elegans* strains used in this study.

| Strain | Genotype | Resource |
| --- | --- | --- |
| NK1316 | *rrf-3(pk1426)* II; *qyls10[lam-1p::lam::GFP]* IV; *rde-1(ne219)* V; *qyls24[cdh-3p::mCherry::PLCδ<sup>PH</sup>]*; *qyls102[fos-1ap::rde-1, myo-2p::GFP]* | Morrissey et al., 2014 |
| NK2144 | *qy17[zmp-1::mNG]* III | Kelley et al., 2019 |
| NK3030 | *qy229[cdh-3p::lmp-1::mNG]* I | This study |
| NK2864 | *qy180[lin-29p::GFP::CAAX]* I | This study |
| NK1173 | *qyls50[cdh-3p::mCherry::moeABD]*; *qyls221[cdh-3p::GFP::ced-10]* | Hagedorn et al., 2013 |
| NK1195 | *qyls127[lam-1p::lam-1::mCherry]* V; *qyls220[cdh-3p::GFP::mig-2]* | Hagedorn et al., 2013 |
| LP399 | *cp151[mNG-C1::3xFLAG::rap-1]* IV | Dickinson et al., 2017 |
| NK3080 | *qy94[mNG::sbp-1]* III; *cpls91[lag-2p::2x mKate2::PLCδ<sup>PH</sup>::3xHA::tbb-2 3'UTR LoxN]* II | This study; Gordon et al., 2019 |
| DG4324 | *tn1765[GFP::3xflag::pod-2]* II | Starich et al., 2020 |
| NK2636 | *qy98[fasn-1::mNG]* I | This study |
| NK2633 | *qy97[elo-1::mNG]* IV | This study |
| NK2950 | *qy208[mNG::lpin-1]* V | This study |
| NK2925 | *qy198[sms-1::mNG]* IV | This study |
| NK2962 | *rrf-3(pk1426)* II; *qy17[zmp-1::mNG]* III | Costa et al., 2023 |
| NK957 | *qyls127[lam-1p::lam-1::mCherry]* V; *qyls66 [cdh-3p::unc-40::GFP]* X; *rol-6(su1006)* II | This study; Hagedorn et al., 2013 |
| NK2935 | *qy202[mNG::hmgr-1]* III | This study |
| NK3017 | *qy215[mNG::fce-1]* I | This study |
| NK2949 | *qy205[lin-29p::icmt-1::mscarlet]* I | This study |
| NK2952 | *qy202[mNG::hmgr-1]* III; *qy205[lin-29p::icmt-1::mscarlet]* I | This study |
| NK3062 | *qy215[mNG::fce-1]* I; *qy239[cdh-3p::icmt-1::mscarlet]* I | This study |
| NK3065 | *qy205[lin-29p::icmt-1::mscarlet]* I; *qy234[mNG::sec16A.1]* III | This study |
| NK3066 | *rrf-3(pk1426)* II; *qyls221[cdh-3p::GFP::ced-10]* | This study |
| NK3055 | *qy234[mNG::sec16A.1]* III; *qy147[lin-29p::mKate2::PLCδ<sup>PH</sup>]* I | This study |
| NK2752 | *qyls555[cdh-3p::aman-2::GFP]*; integrant of *qyEx73[cdh-3p::aman-2::GFP]* | This study; Naegeli et al., 2017 |
| NK2943 | *unc-6(ev400)* X; *qy202[mNG::hmgr-1]* III | This study |
| NK2966 | *unc-6(ev400)* X; *qy205[lin-29p::icmt-1::mscarlet]* I | This study |
| NK2929 | *qy199[fnta-1::mNG]* IV | This study |
| NK2959 | *rrf-3(pk1426)* II; *qy198[sms-1::mNG]* IV | This study |

**Table S4.**
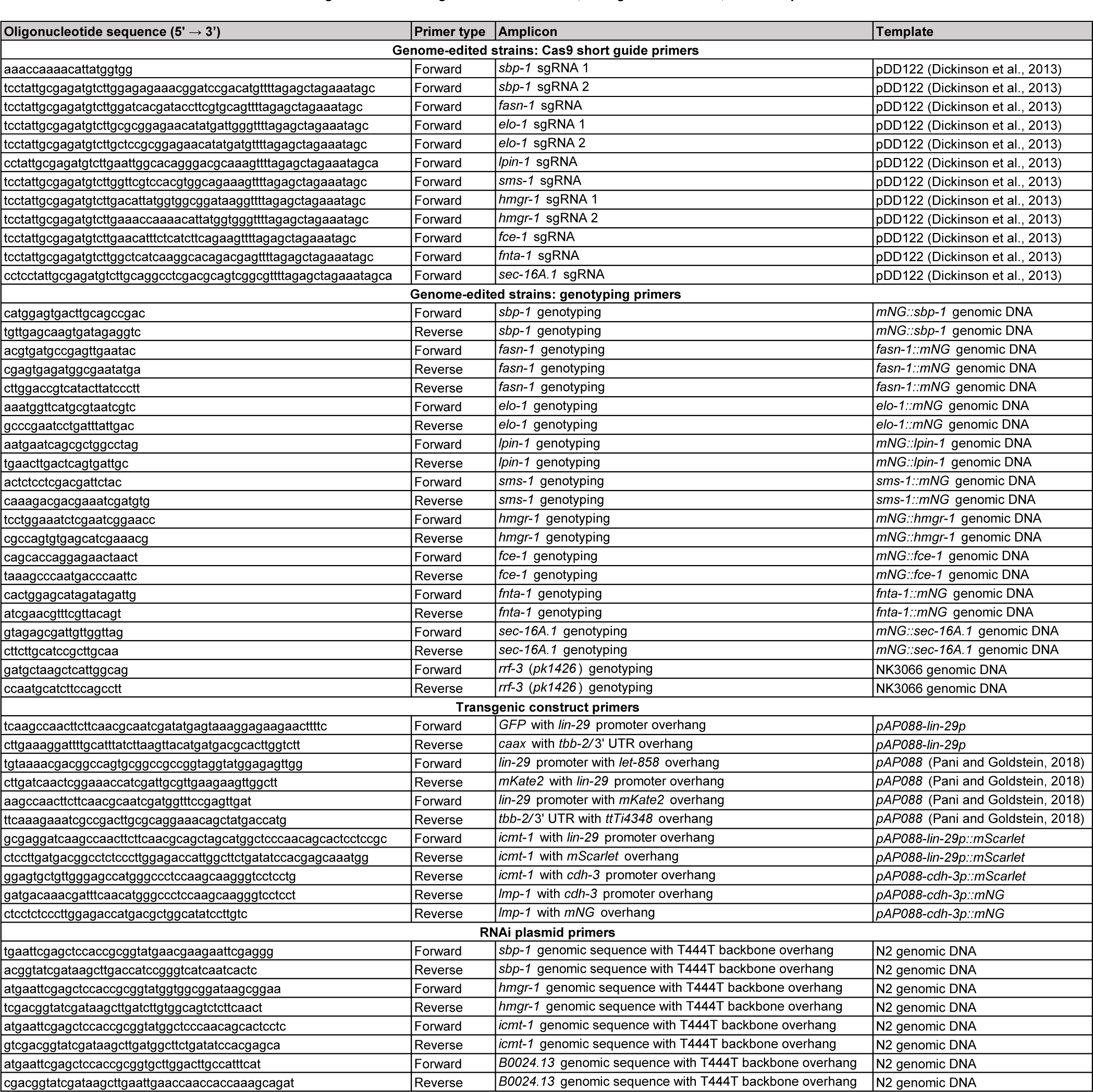
Oligonucleotides for genome-edited strains, transgenic constructs, and RNAi plasmids.

**Table S5.** Chemical and DNA reagents used in this study.

| Reagent | Source/reference | Identifier |
| --- | --- | --- |
| Ampicillin | Sigma-Aldrich | #A0166 |
| Isopropylisopropyl β-D-1-thiogalactopyranoside | Sigma-Aldrich | #I6758 |
| Hygromycin B | Sigma-Aldrich | #H3274 |
| Sodium azide | Sigma-Aldrich | #S2002 |
| Levamisole | Sigma-Aldrich | #L9756 |
| pDD122 (plasmid) | Dickinson et al., 2013 | Addgene_47550 (BRID) |
| pCFJ352 (plasmid) | Addgene | Addgene_30539 (BRID) |
| pAP088 (plasmid) | Pani and Goldstein, 2018 |  |

**Video 1. SBP-1 is required for AC invasive protrusion formation.** Lateral view timelapse images showing the AC invasive protrusion visualized with the membrane marker mCherry::PLCδ^PH^ (cyan) in a control and AC-specific *sbp-1* RNAi knockdown animal. BM was visualized with LAM-1:GFP (magneta). Movies were acquired with a CSU-10 spinning disc confocal microscope. Gaussian blur (1 pixel) was applied to both control and RNAi-treated animals to facilitate visualization of AC protrusion. The total video duration was 75 min, frame interval 5 min, display rate 8 frames/second. Scale bar, 5µm.

**Video 2. LPIN-1 is required for AC invasive protrusion formation.** Lateral view timelapse images showing the AC invasive protrusion visualized with the membrane marker mCherry::PLCδ^PH^ (cyan) in a control and AC-specific *lpin-1* RNAi knockdown animal. BM was visualized with LAM-1:GFP (magneta). Movies were acquired with a CSU-10 spinning disc confocal microscope. Gaussian blur (1 pixel) was applied to both control and RNAi-treated animals to facilitate visualization of AC protrusion. The total video duration was 75 min, frame interval 5 min, display rate 8 frames/second. Scale bar, 5µm.

**Video 3. SMS-1 is required for the AC invasive protrusion formation.** Lateral view timelapse images showing the AC invasive protrusion visualized with the membrane marker mCherry::PLCδ^PH^ (cyan) in a control and AC-specific *sms-1* RNAi knockdown animal. BM was visualized with LAM-1:GFP (magneta). Movies were acquired with a CSU-10 spinning disc confocal microscope. Gaussian blur (1 pixel) was applied to both control and RNAi-treated animals to facilitate visualization of AC protrusion. The total video duration was 60 min, frame interval 4 min, display rate 8 frames/second. Scale bar, 5µm.

**Video 4. HMGR-1 is required for the AC invasive protrusion formation** Lateral view timelapse images showing the AC invasive protrusion visualized with the membrane marker mCherry::PLCδ^PH^ (cyan) in a control and AC-specific *hmgr-1* RNAi knockdown animal. BM was visualized with LAM-1:GFP (magneta). Movies were acquired with a CSU-10 spinning disc confocal microscope. Gaussian blur (1 pixel) was applied to both control and RNAi-treated animals to facilitate visualization of AC protrusion. The total video duration was 75 min, frame interval 5 min, display rate 8 frames/second. Scale bar, 5µm.

